# Phylogenetic Mosaic of an Arms Race with Asymmetrical Sexual Conflict and Its Macroevolutionary Consequences in a Lineage of Small Water Striders

**DOI:** 10.64898/2026.06.24.734260

**Authors:** Zihe Li, Huanyu Chen, Zezhong Jin, Hendrik Freitag, Christine Hecher, Herbert Zettel, Siying Fu, Chen Liu, Mu Qiao, Boxiong Guo, Wenjun Bu, Zhen Ye

**Affiliations:** Institute of Entomology, College of Life Sciences, Nankai University, Tianjin, 300071, China; Museum für Naturkunde, Leibniz Institute for Evolution and Biodiversity Science, Invalidenstraße 43, Berlin, 10115, Germany; 2nd Zoological Department, Natural History Museum Vienna, Vienna 1010, Austria

**Keywords:** Phylogeny, *Pseudovelia*, Sexual conflict, Sexual asymmetry, Speciation, Macroevolution

## Abstract

Sexual conflict has been hypothesized as a driver of speciation, though its effects are likely heterogeneous across phylogenies and between sexes. The Gerromorpha (semi-aquatic bugs), which inhabits water surfaces across diverse aquatic environments, long served as a model for studying sexual conflict. While previous studies have focused on rapid antagonistic coevolution and the genetic basis of sexually antagonistic traits, the macroevolutionary consequences of asymmetrical sexual conflict—particularly male-dominated grasping traits versus female resistance—remain largely unexplored. Within the subgenus *Pseudovelia*, males exhibit pronounced phenotypic diversification in grasping structures, whereas females show modest, clade-specific resistance traits, suggesting male-biased asymmetric conflict. This system presents a valuable opportunity to examine how sexual conflict influences diversification and asymmetrical trait evolution across lineages. Using 204 individuals, representing over half of the subgenus’s species diversity, we reconstructed a time-calibrated phylogeny, quantified diversification rates, assessed sexual conflict intensity across clades, and analyzed correlations between sexual trait evolution and diversification. Our results reveal extensive phylogenetic conflict, particularly within the East Asian clade, driven by introgression and incomplete lineage sorting (ILS). Furthermore, we observe significant phylogenetic heterogeneity in both phenotypic evolution and diversification rates. Notably, a male “trait package” enhancing grasping ability likely drives rapid diversification in the recently radiated “South China” lineage. In contrast, grasping traits involving abdominal segment VIII are associated with lower conflict intensity, facilitating greater evolutionary flexibility in female resistance and resulting in lineage-specific counter-adaptations. These findings highlight the heterogeneous dynamics of asymmetrical sexual conflict in shaping diversification and speciation.

## Introduction

Sexual conflict, which typically arises due to the unequal reproductive payoffs between males and females (Parker, 1979), has long been a central topic in evolutionary biology. In recent years, several theories have proposed that sexual conflict acts as a driver of speciation (Gavrilets, 2014; Mendelson & Safran, 2021). Early studies provided strong support for this view, demonstrating that sexual conflict can promote rapid genetic divergence and reproductive isolation through sexually antagonistic coevolution (Martin & Hosken, 2003; Rice, 1996; Ronkainen et al., 2005). Divergent reproductive interests may also lead to the evolution of pre- and post-mating barriers, such as differences in mating preferences or reproductive organ morphology, thereby facilitating speciation (Gavrilets & Hayashi, 2005; Rice, 1998).

However, growing evidence suggests that the evolutionary consequences of sexual conflict often exhibit substantial mosaic evolution (Carvalho et al., 2019; Crumière et al., 2019; Genevcius et al., 2020; Miller, 2003). A common pattern is that such heterogeneity is clade-specific: sexual conflict traits may not escalate continuously but instead evolve recurrently or via distinct trajectories within a lineage. For instance, *Pteronymia* butterflies have independently evolved male mating plugs (sphragis) multiple times, highlighting clade-specific heterogeneity across the phylogeny(Carvalho et al., 2019). Moreover, the variation in sexual conflict can be not only clade-specific but also sexually asymmetric. The distribution of sexually modified traits may not align strictly between sexes across lineages, with one sex may exhibit delayed or mosaic patterns of trait evolution. A male-biased sexual conflict is evident in diving beetles (Dytiscidae): males evolved adhesive foreleg suction cups once, whereas females subsequently developed resistance traits on the dorsal surface multiple times, nested within the lineage (Miller, 2003). Similarly, in the genus *Rhagovelia* (Veliidae), males have evolved increasingly elaborate grasping structures, yet corresponding female resistance traits have only emerged in clades where males are most “armed”, indicating male-biased escalation (Crumière et al., 2019). Conversely, studies across Coleoptera have shown that evolutionary transitions in genital coevolution often involve changes in female genital morphology preceding those in males, indicating a female-biased pattern (Genevcius et al., 2020). These examples underscore that the tempo, mode and directionality of sexual conflict trait evolution are not universally consistent but are profoundly shaped by linage-specific histories. Critically, two aspects of mosaic evolution remain underexplored, largely because sexual conflict models often examine secondary sexual traits at the level of single species or populations—rarely within a broader phylogenetic context. These aspects are: 1) Clade-specific heterogeneity: whether the intensity of sexual conflict and its impact on diversification rates vary significantly among subclades within a lineage; 2) Sexual asymmetry in trait evolution: how male offensive traits and female resistance traits evolve at different lineages, exhibit divergent morphologies, and follow different phylogenetic trajectories.

The Infraorder Gerromorpha (Insecta: Hemiptera: Heteroptera), commonly known as semi-aquatic bugs, inhabit the water surfaces of diverse aquatic environments and have long served as a model system for studying sexual conflict. In most species within this group, males tend to mate repeatedly, whereas females incur fitness costs from multiple mating or polyandry (Ronkainen et al., 2010), such as increased predation risk and reduced mobility (Arnqvist, 1989; Fairbairn, 1993; Rowe, 1994). This sexual conflict typically drives antagonistic coevolution, with males evolving specialized grasping structures and females developing resistance traits in response (Arnqvist & Rowe, 2002; Crumière et al., 2019; Khila et al., 2012). Although recent work has focused on the genetic basis of such traits (Crumière et al., 2019; Khila et al., 2012), the macroevolutionary consequences of sexual conflict traits across the lineage remain largely unexplored.

Within Gerromorpha, the subgenus *Pseudovelia —* a lineage of small water striders belonging to the genus *Pseudovelia*, subfamily Microveliinae of the family Veliidae, comprises 82 species distributed across the Palearctic, Oriental and Afrotropical regions. This lineage exhibits particularly high diversity in East and Southeast Asia, where 69 species and 2 subspecies have been recorded (Andersen, 1983; Hecher, 2022; Hecher & Laciny, 2021; Li et al., 2022; Linnavuori, 1977; Nieser, 1995; Watanabe & Hayashi, 2023). Despite well-established species classifications, the phylogenetic relationships within *Pseudovelia* remain unclear. Mitochondrial data have revealed non-monophyly among rapidly radiating species (Ye et al., 2016), a phenomenon commonly observed in animals (McKay & Zink, 2010; Noguerales & Emerson, 2025; Ross, 2014). The mitochondrial non-monophyly may result from multiple biological factors, including introgression, ILS, and mitochondrial capture (McKay & Zink, 2010; Noguerales & Emerson, 2025; Ross, 2014). Therefore, a more robust nuclear-based phylogeny, along with investigations into introgression and ILS, is essential for understanding this rapidly radiating subgenus. Males of *Pseudovelia* exhibit remarkable diversity in grasping traits, while females show corresponding counter-adaptations (Figs. 1a–g). The most striking male trait is the highly diversified male abdominal segment VIII (ASE). Most *Pseudovelia* males have elaborate modifications on the ventral side of ASE, including specialized setae, spines or lobe-like processes (Hecher, 2006; Hecher, 2022; Ye et al., 2013), which are also crucial for species identification (Figs. 1b and 1e; Supplementary Fig. S1). Male ASE modifications function to open the female proctiger during copulation were demonstrated in other Microveliinae taxa (Maroni et al., 2023). Like other gerromorphans, female *Pseudovelia* possess concealed genitalia (Han & Jablonski, 2009), which likely necessitates such grasping structures for successful copulation. We therefore consider ASE modifications as putative grasping structures pending future experimental validation. In addition to ASE modifications, some *Pseudovelia* species, particularly East Asian species that likely diverged more recently, also display additional male putative grasping traits, such as tibial processes on the forelegs, strongly curved hind tibiae, and spine-like setae on the hind tibiae (Ye & Bu, 2015; Ye et al., 2016; Ye et al., 2013) (Figs. 1c and d). In Gerromorpha, functional studies have demonstrated that male legs modification serve to grasp and secure females during mating struggles (Arnqvist & Rowe, 2002; Crumière et al., 2019; Khila et al., 2012). Based on these documents and our observations, we infer that the leg modifications in *Pseudovelia* likely serve a similar grasping function. In contrast, female putative resistance traits show relatively modest morphological changes, and morphological studies suggest that different lineages may exhibit distinct phenotypic strategies—such as elongated abdominal tergites (Fig. 1f), modified setae on the gonocoxa (Fig. 1g), and reinforced hind femora (Hecher, 2006; Li et al., 2022), in which the elongated abdominal tergites are documented as female resistance traits in Gerridae (Pruvôt et al., 2024). This pattern reflects a male-biased asymmetric conflict model, which deviates from classical symmetric arms-race models. Therefore, *Pseudovelia* represents an ideal system to investigate how sexual conflict drives diversification, and how sexual asymmetry influences trait evolution across lineages.

**Figure 1.**
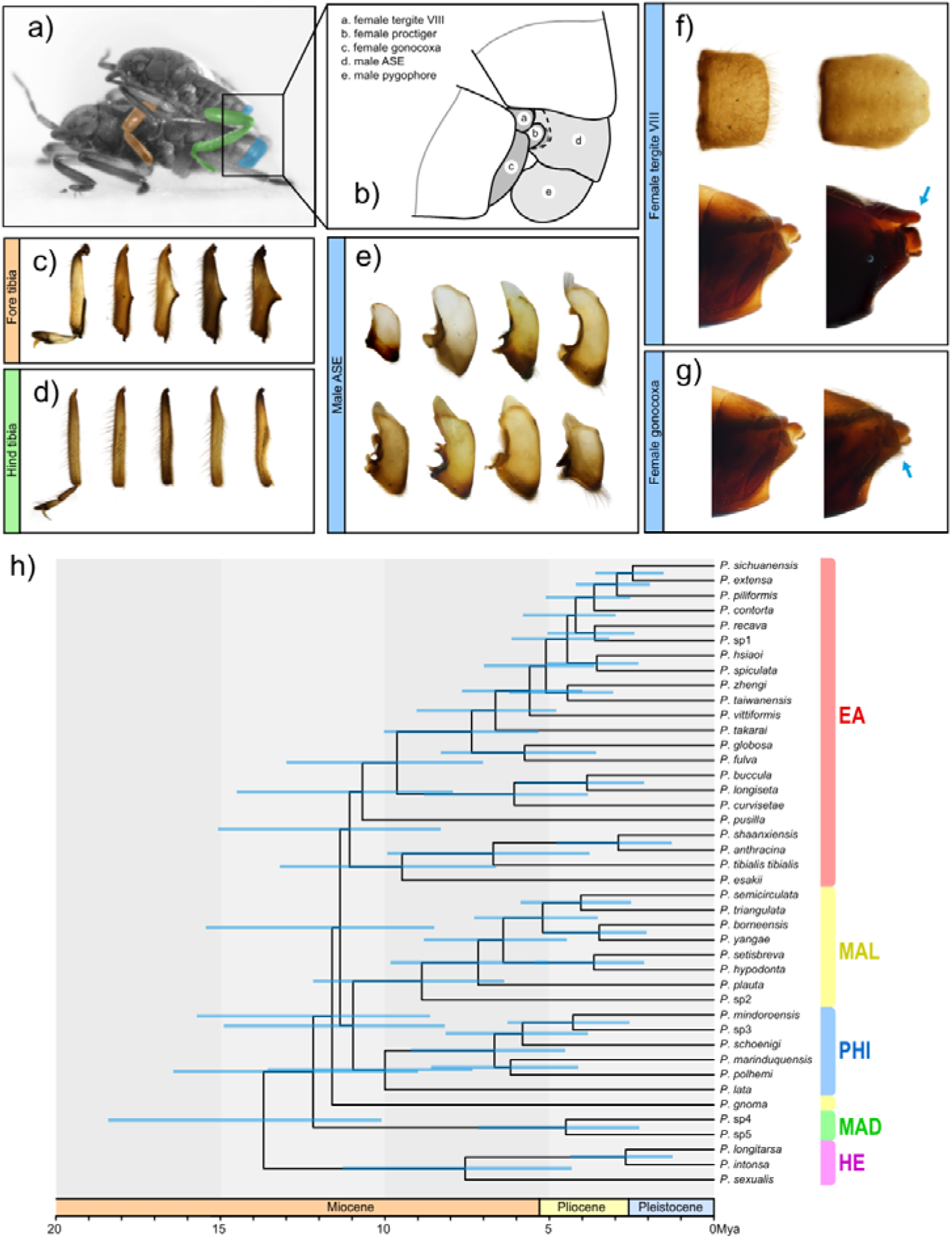
Time-calibrated phylogeny and sexual traits of subgenus *Pseudovelia*. a) – g) Sexual traits of *Pseudovelia*. Photographs and illustrations display male and female traits. a) Photograph of mating pair. b) Illustration of mating pattern. c) Phenotypic disparity in male fore tibiae. d) Phenotypic disparity in male hind tibiae. e) Phenotypic disparity in male abdominal segment VIII (ASE). f) Female tergite VIII: normal (left) and elongated (right). g) Female gonocoxa: normal (left) and with modified setae (right). h) Time-calibrated phylogeny of *Pseudovelia*. Tree topology was inferred using USCOs with ASTRAL, and divergence times were estimated in MCMCtree (PAML) based on fossil-calibrated outgroups. Outgroups are not shown. Node bars represent the 95% highest posterior density (HPD) of divergence times.

In this study, we sampled 204 individuals representing 42 *Pseudovelia* species, accounting for 51.2% of all described species within the subgenus. We reconstructed a robust phylogeny based on whole-genome single nucleotide polymorphisms (SNPs), universal single-copy orthologs (USCOs), and mitochondrial genome (MT), explicitly considering potential confounding factors such as introgression and incomplete lineage sorting (ILS). Furthermore, we conducted fossil-calibrated divergence time estimations and ancestral range reconstructions. Within this framework, we quantified differences in diversification rate among clades and assessed the intensity of sexual conflict in both sexes. We also reconstructed the evolutionary history of key conflict-related traits and analyzed the relationships between trait evolution and diversification. We hypothesize that (1) The multiple male grasping traits, particularly modifications on legs, have evolved into an integrated functional “trait package” in recently diverged East Asian species. This integrated package is expected to intensify sexual conflict and promote lineage diversification; (2) In lineages where male grasping relies primarily on ASE modifications rather than on the highly modified leg structures seen in East Asian species, the intensity of sexual conflict may have remained relatively low, thus allowing evolutionary flexibility in female resistance. This flexibility likely facilitates lineage-specific counter-adaptations and produces a mosaic pattern of female resistance traits across subclades.

## Materials & Methods

Our workflow is summarized in Figure 2.

**Figure 2.**
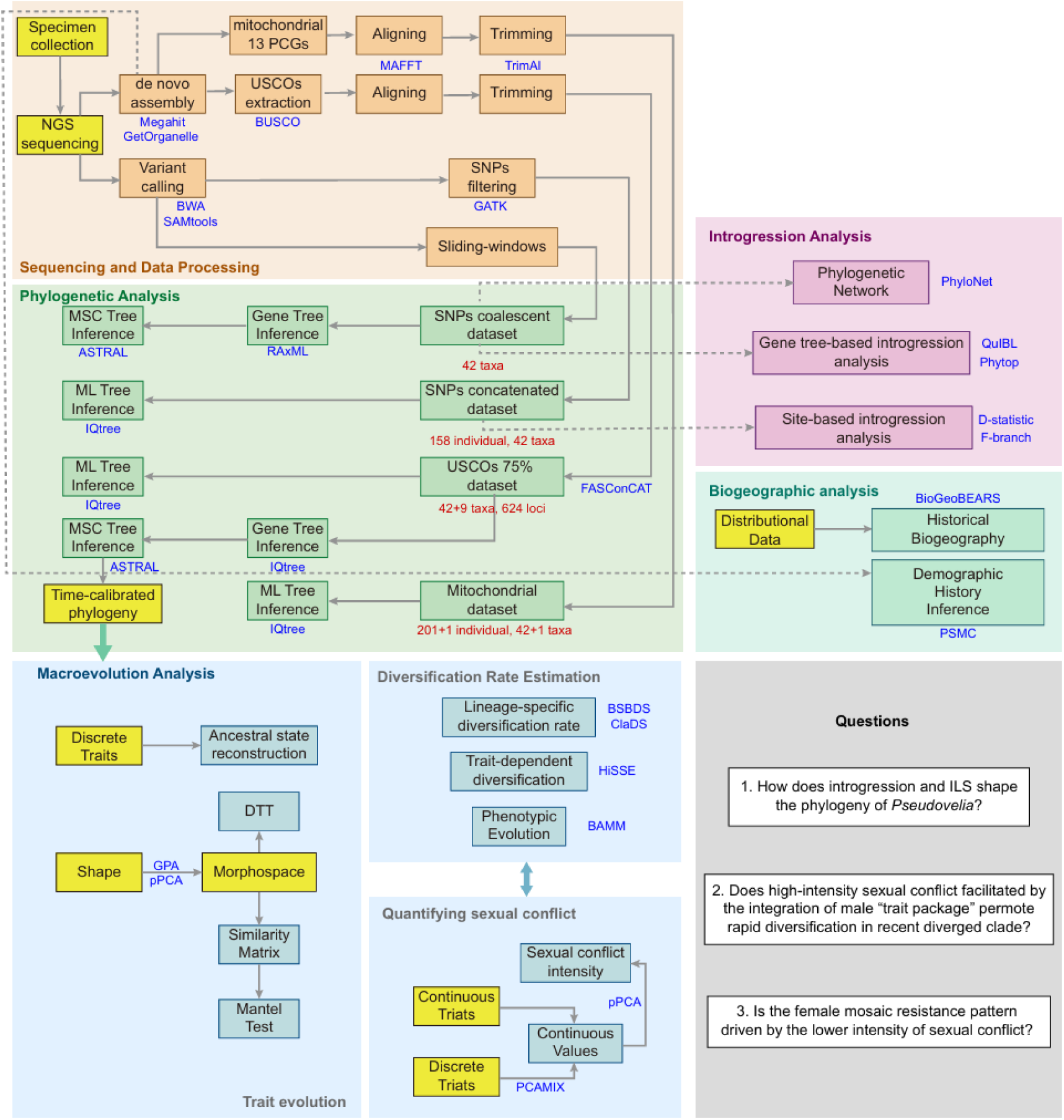
Analytical pipeline of this study.

### High-Quality Reference Genome for P. tibialis tibialis

#### DNA extraction

Specimens of *P. tibialis tibialis* used for reference genome sequencing were collected from Xianning city, Hubei Province, China. Genomic DNA was extracted using the SDS method, followed by purification with QIAGEN Genomic kit (Cat#1343, QIAGEN), according to the manufacturer’s standard operating protocol. DNA integrity was assessed via 1% agarose gels electrophoresis. Purity was evaluated using a NanoDrop One UV-Vis spectrophotometer (Thermo Fisher Scientific, USA), and DNA concentration was measured with a Qubit 3.0 Fluorometer (Invitrogen, USA).

#### Library preparing and sequencing

For Oxford Nanopore Technologies (ONT) sequencing, 2 μg of high-quality genomic DNA was used as input material for library preparation. After quality assessment, long DNA fragments were size-selected using the BluePippin system (Sage Science, USA). End-repair and A-tailing were performed using the NEBNext Ultra II End Repair/dA-tailing Kit (Cat#E7546). Adapter ligation was carried out with the LSK109 kit (Oxford Nanopore Technologies, UK). The library fragment size was confirmed using a Qubit 3.0 Fluorometer (Invitrogen, USA). Sequencing was subsequently conducted on a Nanopore GridION X5/PromethION sequencer (Oxford Nanopore Technologies, UK) at Nextomics (Wick et al. 2019).

#### Data quality control

Nanopore sequencing generated FAST5 files containing raw signal data.

Basecalling was performed using Guppy v3.2.2+9fe0a78 (Sherathiya et al., 2021) to convert the FAST5 files into FASTQ format. Raw FASTQ reads with a mean_qscore_template < 7 were filtered, and only pass reads were retained.

#### De novo assembly

De novo assembly was performed using an ONT-only approach based on the string graph method implemented in NextDenovo v2.3.1 (https://github.com/Nextomics/NextDenovo.git). Due to the high error rate of ONT raw reads, initial subreads were self-corrected using the NextCorrect module to generate consistent sequences (CNS reads). These CNS reads were then processed with the NextGraph module to identify their correlations, which were used to assemble the preliminary genome. To improve assembly accuracy, the contigs were polished using Racon with ONT long reads and NextPolish v1.3.0 (Hu et al., 2020) with Illumina short reads under default settings. Potentially redundant contigs were removed by conducting similarity searches with the parameters “identity 0.8 –overlap 0.8”.

Genome assembly completeness was evaluated using BUSCO v4.0.5 (Benchmarking Universal Single Copy Orthologs) (Simão et al., 2015) and CEGMA (Core Eukaryotic Gene Mapping Approach) (Parra et al., 2007). To assess assembly accuracy, Illumina paired-end reads were mapped to the assembly using BWA (Burrows-Wheeler Aligner) (Vasimuddin et al., 2019), and mapping rate along with genome coverage was calculated using SAMtools v0.1.1855 (Danecek et al., 2021). Base-level accuracy was further assessed using BCFtools v1.8.0 (Li & Durbin, 2011).

To evaluate gene coverage, RNA-seq reads were aligned to the assembled genome using HISAT (Kim et al., 2015) with default parameters. To exclude mitochondrial sequences, the draft genome was aligned to the NCBI NT database, and matching sequences were removed from the assembly.

#### Taxon Sampling and Resequencing

We sampled 204 newly sequenced individuals representing 42 *Pseudovelia* species (Supplementary Table S1). Additionally, we included one individual each from the following outgroup species for resequencing: *Microvelia horvathi*, *Baptista hoedli*, *Aquarius paludum*, *Limnoporus rufoscutellatus*, *Cylindrostethus scrutator*, *Perittopus yunnanensis*, *Rhagovelia sumatrensis*, *Entomovelia doveri* and *Entomovelia quadripenicillata*. All specimens were presented in absolute alcohol and stored at –20 . Genome DNA was extracted using CWBIO Universal Genomic DNA Kit and subsequently stored at –80 . Whole-genome resequencing was performed using 150 bp pair-ends reads on the NovaSeq 6000 platform (Biomarker Technologies, Beijing, China).

#### Variant Calling

Sequencing reads were mapped to the reference genome using BWA v0.7.18 (Li & Durbin, 2009). Resulting BAM files were sorted, and duplicates removed using SAMtools v1.7 (Li et al., 2009). Mapping and coverage statistics were computed using the flagstat and depth modules of SAMtools. SNP calling and filtering were conducted using GATK v4.1.9.0 (McKenna et al., 2010). Insertions and deletions (INDELs) were excluded via the GATK SelectVariants module. SNPs were retained based on the following criteria: minimum quality value ≥ 30 (--minQ 30); minimum mean deapth ≥ 5 (--min-meanDP 5); retention of only biallelic sites (--min-alleles 2, --max-alleles 2), and exclusion of sites with missing rate > 0% (--max-missing 1). The final filtered VCF file was converted to FASTA format using vcf_tab_to_fasta_alignment.pl (https://github.com/JinfengChen/vcf-tab-to-fasta).

### Phylogenetic Analysis

#### Mitochondrial dataset

A total of 201 ingroup individuals (42 species) and one outgroup (*Microvelia douglasi*) were included. Complete mitochondrial genomes were assembled from resequencing reads using GetOrganelle v1.6.4 (Jin et al., 2020) with the animal_mt dataset and k-mer sizes of 21, 45, 65, 85 and 105. The 13 protein-coding genes (PCGs) were extracted for phylogenic analysis. Alignment, trimming, and concatenating of PCGs were performed in Phylosuite v1.1.16 (Zhang et al., 2020). A maximum likelihood (ML) tree was reconstructed using IQtree2 (Nguyen et al., 2015) under the ModelFinder Plus (MFP) mode with 1,000 ultrafast bootstrap replicates for branch support.

#### SNP concatenated dataset

To reduce computational burden while maintaining taxonomic coverage, we subsampled the SNP dataset to 158 individuals by retaining only one or two high-coverage individuals per population. A concatenated SNP matrix of the 158 individuals from 42 species was used to build an ML tree in IQtree2 (MFP mode; 1,000 bootstrap replicates). A multi-species coalescent tree was also inferred using SVDquartets (Chifman & Kubatko, 2014) in PAUP* v4.0a (Swofford, 2000) under the “motilocus” model with 100 bootstrap replicates and the “mscoalescent” tree model, seeded at 1100.

#### Universal single-copy orthologues (USCOs) species tree

One individual per species (42 ingroup + 9 outgroup) was selected. Genome contigs were assembled with Megahit v1.2.9 (Li et al., 2016), and USCOs indentified via BUSCO v5 (Simão et al., 2015) using the hemiptera odb_10 dataset. Nucleotide sequences were aligned using MAFFT (Nakamura et al., 2018), trimmed with trimAl (Capella-Gutierrez et al., 2009), and concatenated using FASConCAT (Kuck & Meusemann, 2010) to produce a 75% completeness matrix. All steps followed scripts from Du et al. (2023). For the concatenated dataset, an ML tree was constructed in IQ-TREE2 with MFP and 1,000 bootstraps. For the coalescent approach, individual gene trees (from the 75% matrix) were built using IQ-TREE2 with automatic model selection, and a species tree was inferred with ASTRAL (Zhang et al., 2018).

#### SNP sliding-window species tree

To validate relationships inferred from USCOs, we generated a species tree using SNP sliding windows. One high-coverage individual per ingroup species was selected (outgroups excluded due to poor mapping). We used non-overlapping 20 kb windows with a step size of 400 kb to reduce recombination impact. Analyses were conducted across all 42 species and within three major clades separately to assess introgression and incomplete lineage sorting (ILS). Windows with < 500 parsimony-informative sites (PIS) were discarded. ML trees were inferred for each window using RAxML (Stamatakis, 2014) with the GTRGAMMA model and 100 bootstrap replicates. All gene trees were rooted using the basal ingroup species from the USCOs coalescent tree. A coalescent tree was generated from these window trees using ASTRAL.

### Divergent Time Estimating

Divergence times were estimated using MCMCTree in PAML (Yang, 2007) based on the USCOs coalescent species tree. Four fossil-constrained nodes in the outgroup species were used for calibration, with the root age fixed at 120 Ma, following prior analyses (Supplementary Table S3). We implemented an independent rates model (clock = 2) to accommodate branch-specific rate variation and selected the HKY85 nucleotide substitution model (model = 4). The MCMC analysis included a 2,000,000-interation burn-in, followed by sampling every 100 iterations to obtain 100,000 samples.

### Introgression and ILS Inference

We used Dsuite (Malinsky et al., 2021) to detect introgression signals among species. The analysis employed a genomic dataset comprising one individual per species, with *P. sexualis* designated as the outgroup. SNP filtering parameters matched those described in the “Variant Calling” section. First, we applied the Dsuite Dtrios module to detect introgression signals based on SNP data. This method assesses allele frequency patterns across quartets (P1, P2, P3 and the outgroup) to compute Patterson’s D-statistics (ABBA-BABA test). Subsequently, we used the Dsuite Fbranch function in conjunction with the topology of the SNP sliding-window tree to infer phylogenetic relationships and lineage-specific introgression. Fbranch values were estimated to quantify deviations from the species tree topology. To improve the resolution of introgression detection, we performed D-statistics and Fbranch analyses separately for the three major clades using the same protocol. Given the large number of species and the evident heterogeneity of the East Asia clade, we classified the 11 most divergent EA species as the “South China” group, representing a distinct species group for the purposes of our introgression and ILS analyses.

SNP sliding-window trees were used to detect introgression and ILS via gene tree-based methods. First, we used Phytop (Shang et al., 2025) to detect ILS or introgression signals at internal nodes of the species tree. For a given topology, such as ((A,B)C), Phytop quantifies the proportion of incongruent topologies [e.g., ((A,C)B) and ((B,C)A)] inferred from the input ASTRAL tree, and then calculates the proportion of gene trees explained by introgression (denoted as IH) or ILS. The indices for ILS (ILS-I, from 0 to 100%) and introgression (IH-I, from 0 to 50%) reflect signal strength, while ILS-e and IH-e represent the actual proportion of incongruent topologies explained by ILS or introgression, respectively. We considered ILS-i > 50% or IH-i > 30% indicative of strong ILS or introgression signals. Statistical significance (assessed via chi-square tests, P < 0.05) was evaluated against the multispecies coalescent model. Our sliding-window SNP ASTRAL tree was used as input. Furthermore, we conducted QuIBL analysis (Edelman et al., 2019) on the gene trees. QuIBL quantifies introgression and ILS among lineages by estimating the internal branch length of three-species triplets. Major lineages were tested separately, and all possible triplet combinations among the input species were examined. To discern whether introgression or ILS was the primary contributor to gene tree discordance, we calculated the delta BIC score. A delta BIC > 10 supported the ILS-only model, while a delta BIC < –10 favored the ILS + introgression model.

### Phylogenetic Network

We inferred phylogenetic networks using PhyloNet (Wen et al., 2018) on 20-kb sliding-window trees under a maximum pseudo-likelihood (MPL) algorithm. The number of reticulation events was explored from 1 to 3, with five independent searches per reticulation. Edges with bootstrap support < 70% were excluded. Networks were visualized in Dendroscope v3.8.10 (Huson & Scornavacca, 2012).

### Analysis of Trait Evolution

We performed ancestral state reconstruction of male and female sexual traits using the fitMk function in the R package phytools (Revell, 2012). For each trait, three evolutionary models were evaluated: equal-rates (ER), symmetric (SYM), and all-rates-different (ARD). Model comparisons were based on Akaike Information Criterion (AIC) values, and the best-fitting model was used to simulate 1,000 stochastic character maps via the make.simmap function.

To quantify correlations among sexual traits, we digitized the contours of male abdominal segment VIII (ASE) in lateral view, inner margin of fore tibiae, and inner margin of hind tibiae using the digitizeImange function in R package StereoMorph (Olsen & Westneat, 2015). The resulting curve data were compiled into a three-dimensional array, with dimensions corresponding to the number of points per curve, the coordinate dimensions (X and Y), and the number of samples. We applied Generalized Procrustes Analysis (GPA) to align and scale shapes using the gpagen function in the R package geomorph (Adams & Otárola-Castillo, 2013), followed by phylogenetic Principal Component Analysis (pPCA) using Generalized Least Squares (GLS) regression via the gm.prcomp function in the R package geomorph, to identify major axes of shape variation and generate a phylomorphospace. The ancestral state (root node) was estimated from the pPCA model. For each extant taxon, we computed displacement vectors in morphospace by subtracting the principal component (PC1–PC3) coordinates from the ancestral state. Similarity matrices were constructed by calculating vector dot product between species’ PCA scores and the ancestral coordinates. Trait shape correlations were assessed using Mantel tests implemented in the R package vegan (Adams & Otárola-Castillo, 2013) with 999 permutations.

For comparative evolutionary trajectory analysis, Procrustes-aligned coordinates were subjected to GPA as described above and used Principal Component Analysis (PCA) via the gm.prcomp function in the R package geomorph. Disparity-through-time (DTT) analysis was conducted along the phylogeny using PC1 scores via the dtt function in R package geiger (Pennell et al., 2014).

### Quantifying the Intensity of Sexual Conflict

We selected two continuous traits (i.e., male-to-female body length ratio and the proportion of grasping comb on male tibiae) and four discrete traits, i.e., (1) presence/absence and type of process (none, angular-like, or spine-like) on male fore tibiae, (2) curvature of male hind tibiae (curved vs. nearly straight), (3) presence/absence of spine-like setae on male hind tibiae, and (4) complexity of male ASE (complex vs. simple), to quantify the intensity of sexual conflict. Discrete traits were converted to continuous variables using Principal Component Analysis for Mixed Data (PCAMIX). To achieve a better dispersion in the phylomorphospace, we applied PCA to perform a linear transformation on the one-dimensional continuous trait data. These transformed variables were integrated into a phylogenetic Principal Component analysis (pPCA) incorporating branch-length scaling. We extracted the first principal axis (pPC1) from the pPCA results and performed a linear transformation to represent sexual conflict intensity on a standardized scale from 0 to 10. For putative female counter-adaptations, we estimate the relative strength of the hind femur by calculating the ratio of the square of its diameter to the cube of the body length. This value was similarly rescaled (0 to 10) for comparative analysis.

### Net Diversification Rate Estimating

To assess the relationship between sexual conflict and diversification rates, we estimated diversification rates using multiple methods based on our time-calibrated phylogeny. First, we implemented the Branch-Specific Birth-Death-Shift (BSBDS) model in RevBayes (Höhna et al., 2016), running an MCMC analysis for 100,000 generations. For cross-validation, we applied ClaDS2 using the R package PANDA (Morlon et al., 2016) under default parameters.

Phenotypic evolutionary rates were inferred using the “trait” module in Bayesian Analysis of Macroevolutionary Mixtures (BAMM) (Rabosky et al., 2014), treating the intensity of both male and female sexual conflict as continuous traits. Priors were scaled to the data using the setBAMMpriors function in the R package BAMMtools. MCMC simulations were run for 50 million generations, with event data sampled every 10,000 generations.

### Association Between Sexual Conflict and Speciation

We extracted net diversification rates for all terminal taxa from the BSBDS analysis in RevBayes, and conducted a linear regression model (ordinary least squares) to evaluate the relationship between the net diversification rate and expression of sexual traits using the base lm function in R. Additionally, to account for phylogenetic non-independence in net diversification rates, we performed phylogenetic generalized least squares (PGLS) analyses using the gls function in the nlme package (Pinheiro, 2025). The overall model significance was assessed using an F-test. A p-value threshold of 0.05 was used to determine statistical significance. Additionally, we used the Hidden State Speciation and Extinction (HiSSE) model (Beaulieu & O’Meara, 2016) to evaluate the effects of discrete male sexual conflict traits on diversification. Thirteen models were employed to examine the association between four discrete traits (presence/absence of distinct processes on fore tibiae, spine-like setae on hind tibiae, hind tibiae curvature, and ASE complexity) and diversification rates.

### Demographic History Inference

We inferred the demographic history of *Pseudovelia* species using the Pairwise Sequentially Markovian Coalescent (PSMC) method (Li & Durbin, 2011). To address low coverage in distantly related species, individual-specific pseudo-reference genomes were constructed from MEGAHIT-assembled contigs using iTools. Input files for PSMC were generated using the mpileup module of SAMtools, with the parameter “-C” set to 50. PSMC analyses were run with 25 iterations (-N25), a maximum coalescent time of 15 (-t15), and a theta/rho ratio of 5 (-r5). The 64 time intervals were set as “4 + 25 ∗ 2 + 4 + 6”. PSMC plots were scaled using a mutation rate (μ) of 3.5 × 10^-9^ and a generation time (g) of 0.5 years across all species.

### Ancestral Range Estimation

Biogeographic histories were reconstructed using the R package BioGeoBEARS. Four geographic regions were defined: East Asia, Malaysia, the Philippines, and Africa. We compared the DEC and DEC+j models, selecting the best-fit model based on AIC values.

## Results

### Phylogenetic Analysis

Phylogenetic relationships of *Pseudovelia* were reconstructed using whole-genome SNPs, USCOs, and mitochondrial PCGs. The concatenated SNP dataset included 158 ingroup individuals representing 42 species, totaling 9,647,404 bp after filtering. The individual mapping and coverage rates are provided in Supplementary Table S2. The USCO dataset, with 75% completeness, comprised 42 ingroup and 9 outgroup individuals, spanning 759,942 bp across 624 loci. Analysis resolved *Pseudovelia* into five distinct clades: East Asia (EA) clade, Malaysia clade (MAL), the Philippines clade (PHI), Madagascar clade (MAD) and Hairy-eyes clade (HE).

A widespread discordance was observed between the mitochondrial and nuclear gene trees. The ML tree based on the 13 mitochondrial PCGs exhibited pervasive non-monophyly, particularly within the EA clade (Fig. 3a; Supplementary Fig. S2). In contrast, both the ML and SVDquartets trees based on concatenated SNPs recovered all species as monophyletic (Fig. 3a; Supplementary Figs. S3 and S4). Regarding relationships among major clades, the HE clade was sister to the EA clade in the mitochondrial tree, whereas in the USCOs trees, it was sister to all other clades (Fig. 3a; Supplementary Figs. S2 and S6).

**Figure 3.**
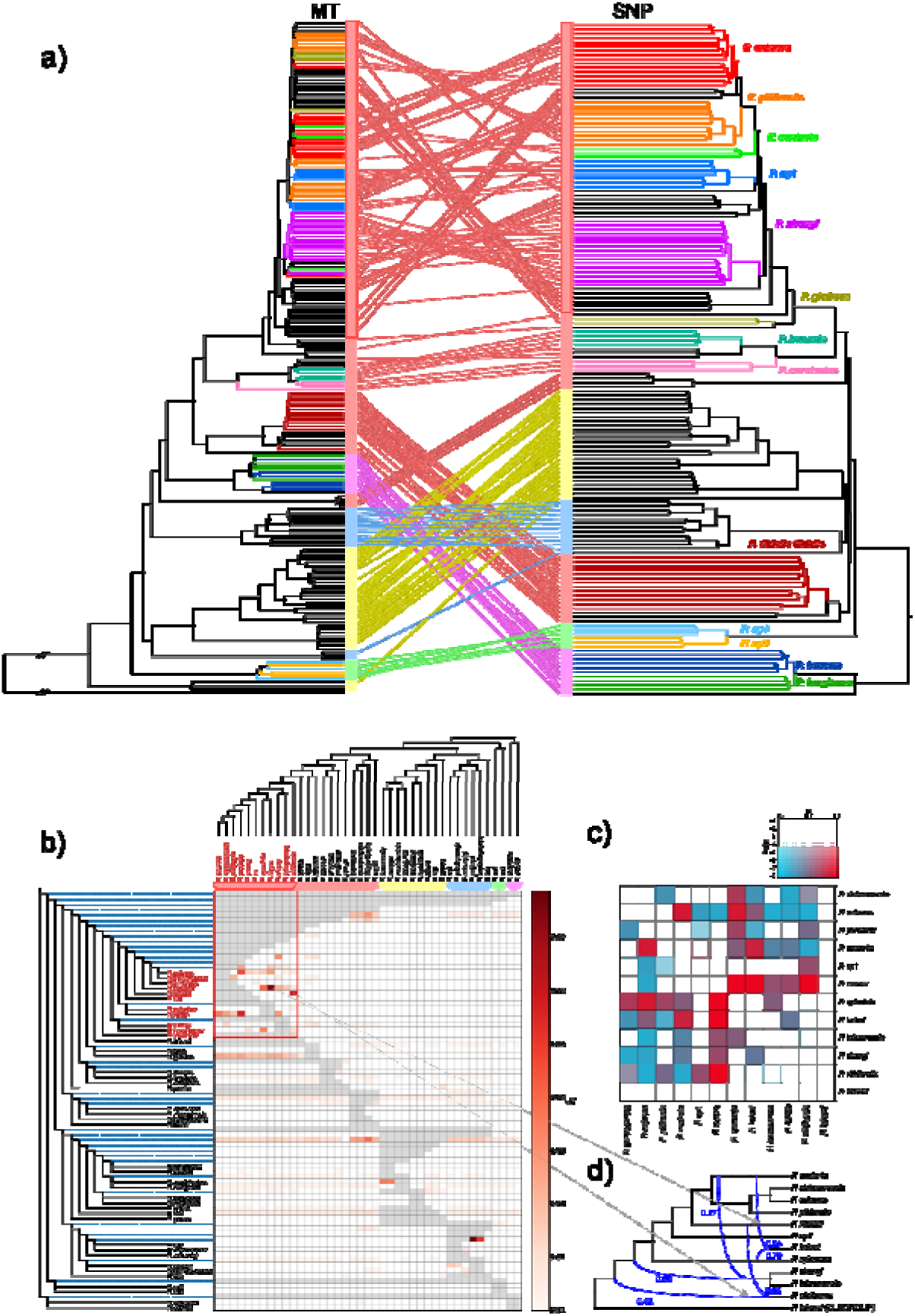
Mito-nuclear discordance and introgression analyses across *Pseudovelia*. a) Phylogenetic trees reconstructed using mitochondrial genomes (left) and SNPs (right) with maximum likelihood (ML) in IQ-tree2. Colored branches indicate non-monophyletic species in the mitochondrial tree. Colored bars next to the trees represent the major clads as identified in Figure 1. b) Matrix of f-branch values, based on the SNP sliding-windows species tree topology, with *P. sexualis* as the outgroup. Values (*f_b_*) were calculated using Dsuite. Colored bars represent the major clads as identified in Figure 1. Species names in red and the matrix enclosed in the red box represent the South China group. c) Matrix of D-statistic values from Dsuite analysis within the South China group. Darker colors indicate smaller *p*-value; red indicates higher D-statistic value, and blue indicates lower values. d) Phylogenetic network analysis in PhyloNet within the South China group (h=3). Blue lines indicate reticulation events; numbers denote inheritance probabilities.

To further investigate topological conflicts, SNP trees were re-rooted using the HE clade as the outgroup, reflecting its basal position in the rooted USCO trees. Comparing USCO and re-rooted SNP trees revealed that ML analysis of both datasets consistently placed *P. shaanxiensis*, *P. anthracina*, *P. tibialis tibalis*, and *P. esakii* as sister to the MAL+PHI clade (Fig. 3a; Supplementary Figs. S3 and S6). However, coalescent-based methods placed these species as sister to the EA clade, and recovered the EA clade as monophyletic (Figs. 1h and 3a; Supplementary Figs. S4–S6). Relationships among species within each major clade were generally stable, except among recently diverged taxa in the EA clade, where persistent conflicts were observed. The placement of *P. gnoma* was notably unstable: SNP trees based on both concatenated and sliding-window datasets consistently placed it as sister to other MAL species (Fig. 3a; Supplementary Figs. S3–S5), whereas the USCO ML tree placed it as sister to *P. pusilla* (Supplementary Fig. S6), and the USCO ASTRAL tree placed it as sister to the EA + MAL + PHI clade (Fig. 1a; Supplementary Fig. S6).

Within the EA clade, the South China group represent the most divergent lineage and exhibited notable phylogenetic conflict among ML and MSC trees inferred from USCOs and SNP datasets (Supplementary Figs. S3–S6). Despite this inconsistency, ASTRAL trees based on USCOs and SNPs showed largely congruent topologies, with the exception of the placement of *P. recava* (Supplementary Figs. S5 and S6). In the USCOs tree, *P. recava* was recovered as sister to *P.* sp1 (Supplementary Fig. S6), whereas in the SNP tree is sister to *P. extensa* + *P. sichuanensis* + *P. piliformis* + *P. contorta*. However, the ancestor node of *P. recava* and *P. extensa* + *P. sichuanensis* + *P. piliformis* + *P. contorta* in SNP tree received very low support (0.02) (Supplementary Fig. S5).

### Divergent Time Estimation

The crown group of *Pseudovelia* originated at 13.72 Ma (95% highest posterior density [HPD]: 10.13–18.45 Ma) (Fig. 1h; Supplementary Fig. S7). Within the genus, the MAL + PHI clade and the EA clade diverged at 11.39 Ma (95% HPD: 8.52–15.48 Ma). The MAL clade originated at 8.90 Ma (95% HPD: 6.37–12.19 Ma), while the PHI clade originated at 10.02 Ma (95% HPD: 7.36–13.60 Ma). The EA clade emerged at 1.10 Ma (95% HPD: 8.30–15.09 Ma), whereas the South China group originated at 5.61 Ma (95% HPD: 4.03–7.67 Ma).

### Introgression and ILS Inference

Our findings revealed significantly stronger and more widespread introgression signals among recently diverged taxa in the South China group compared to other clades (Fig. 3b). Within the PHI clade, significant introgression was also detected between *P. marinduquensis*, *P. polhemi*, and the ancestral node of *P.* sp3 + *P. mindoroensis* (Fig. 3b). Although no significant inter-clade introgression was identified, weak signals were observed between the ancestral node of the MAL clade and most species (Fig. 3b). These patterns were further supported by D-statistics analyses (Supplementary Fig. S8). Independent D-statistic and f-branch analyses conducted within the three major clades were consistent with the genus-wide results (Figs. 3c and 3d; Supplementary Figs. S9–S11).

Phytop analysis quantified the contributions of introgression and ILS at internal nodes. Using 2,907 window trees filtered by PIS to construct the species tree, we identified six internal nodes within the EA clade exhibiting significantly high levels of both ILS (ILS-i > 50%) and introgression (IH-i > 30%, P < 0.05), and one node showing only significant introgression (ILS-i < 50%, IH-i > 30%, P < 0.05). Introgression-only signals were also detected at the ancestral nodes of the MAL clade and the EA + MAL + PHI clade (Supplementary Fig. S12). We also conducted QuIBL analysis separately on the three major clades. For the South China group within the EA clade, 2,997 trees generated from 20-kb windows (filtered by PIS) were analyzed. Results showed that 82.4% of triplets (136/165) supported the ILS model (ΔBIC > 10), whereas only 2.4% (4/165) supported the introgression model (ΔBIC < –10), suggesting that incomplete lineage sorting (ILS) was the primary source of phylogenetic discordance (Supplementary Table S4). In the MAL clade, analysis of 2,941 trees revealed that 51.0% (48/84) supported ILS and 17.9% (15/84) supported introgression (Supplementary Table S5). In contrast, analysis of 2,956 trees in the PHI clade found no support for ILS, while 50.0% (10/20) supported introgression (Supplementary Table S6).

### Phylogenetic Network

PhyloNet analyses were performed with reticulation numbers set from h = 1 to 3. At h = 1 and 2, ancestral gene flow was inferred. At h = 1, gene flow occurred from the ancestor of the MAL clade to both *P. gnoma* and the ancestor of the PHI clade; at h = 2, gene flow extended from the MAL + PHI ancestor to both *P. gnoma* and the MAD clade. Only recent gene flow was detected at h = 3 (Supplementary Fig. S8).

To improve detection sensitivity, each major clade was analyzed independently. For the EA clade’s South China group (with *P. takarai* as the outgroup), gene flow was detected from *P. recava* to *P. vittiformis*, and from *P. hsiaoi* to *P. contorta* and *P. spiculata* at h = 1 and 2—consistent with site-based analyses. However, the reticulation involving *P. contorta* and *P. zhengi* + *P. taiwanensis* detected at h = 3 lacked support in site-based analyses (Supplementary Fig. S9). For the MAL clade, only one reticulation event was found across h = 1 to 3, consistent with site-based results (Supplementary Fig. S10). For the PHI clade, all models detected reticulations involving *P. polhemi* and the ancestral node of *P.* sp3 + *P. mindoroensis* (Supplementary Fig. S11).

### Analysis of Trait Evolution

Model comparison using AIC values indicated that the All-Rates-Different (ARD) model best fitted the data for all sexual traits in our ancestral state reconstruction analysis. The reconstruction of six sexual traits revealed that these traits remained unchanged in the common ancestor of the subgenus *Pseudovelia* (Supplementary Fig. S13). Male leg modifications, however, were restricted to recently diverged species within the EA clade (Fig. 4a; Supplementary Fig. S13). The male fore tibial process and spine-like tibial setae each evolved independently once, while hind tibial curvature evolved independently twice. Both the fore tibial process and hind tibial curvature were secondarily lost in *P. sichuanensis* (Supplementary Fig. S13). All transitions from simple to modified male leg traits occurred around 5.11 Ma (95% HPD: 3.64–7.00 Ma), coinciding with recent speciation events and the peak of phenotypic disparity (Fig. 4a, b and c; Supplementary Fig. S13). In contrast, the evolution of complexity in the male ASE occurred once at the ancestral node of the EA+MAL+PHI clade, and was subsequently lost in *P. buccula*, *P. longiseta*, *P. curvisetae* and *P. pusilla* (Fig. 4a; Supplementary Fig. S13). Among female resistance traits, elongated tergite VIII and modified gonocoxal setae each evolved once within the MAL and PHI clades, respectively (Fig. 4a; Supplementary Fig. S13). The reinforced female hind femur was characterized by a sexual conflict intensity value greater than 5, and this reinforcement evolved independently in *P. sichuanensis*, *P. contorta*, *P. spiculata* and *P.* sp4, all within the EA clade, with the exception of *P*. sp4 (Fig. 4a; Supplementary Fig. S13). Overall, the distribution of sexual traits suggests a clade-specific pattern of sexual conflict in *Pseudovelia*.

**Figure 4.**
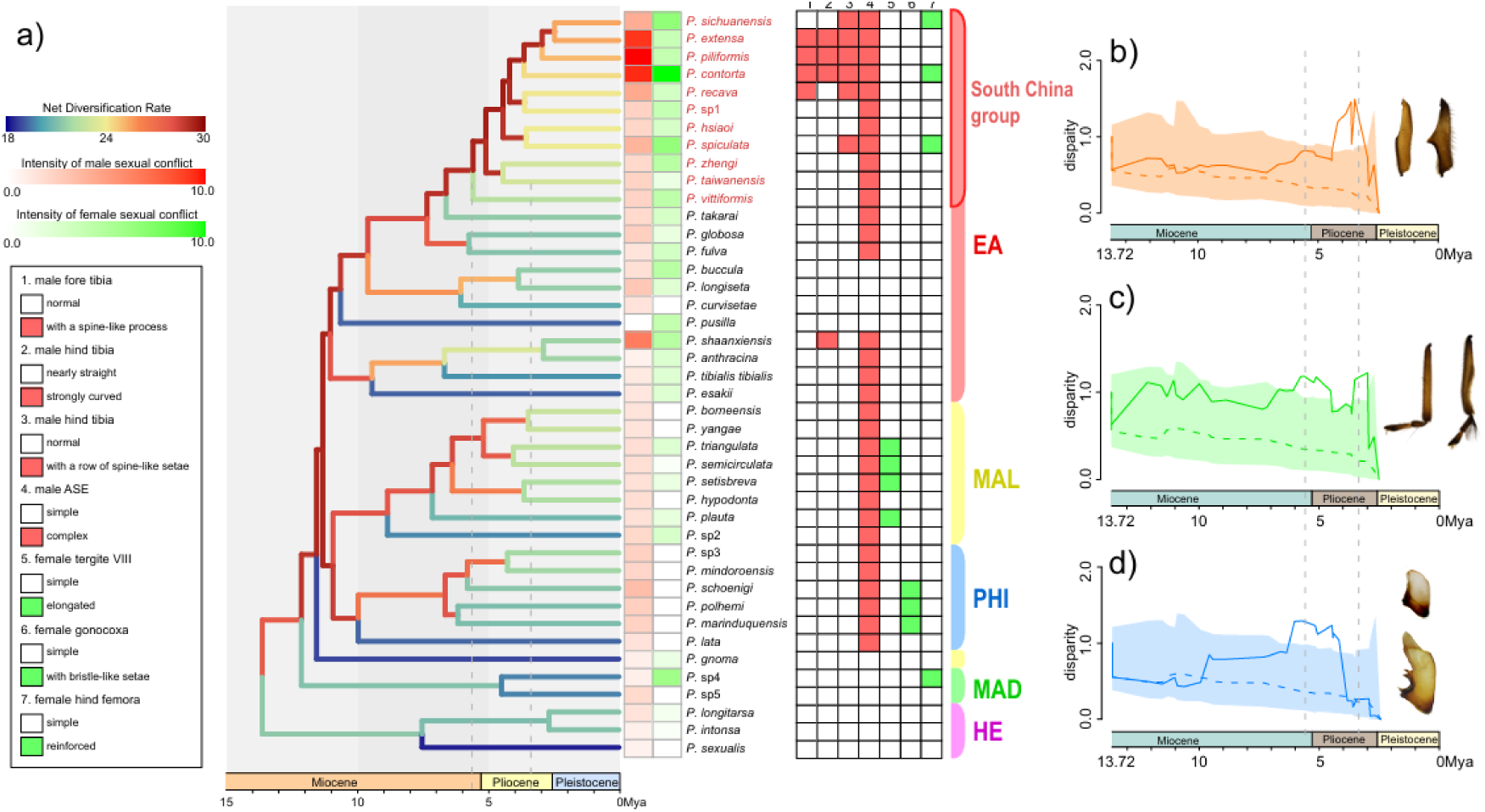
Macroevolution of sexual conflict in *Pseudovelia*. a) Net diversification rate, expression of sexual traits, and ancestral state reconstruction across lineage. Branch colors represent lineage-specific net diversification rates inferred by RevBayes. Dotted lines mark the time of peak phenotypic disparity in disparity-through-time (DTT) analysis. Red bars indicate expression of male traits, and green bars indicate expression of female traits. States of sexual traits are showed on the right; red squares represent modified male traits, and green squares represent modified female traits. Colored bars on the right represent the major clads as identified in Figure 1. Species names in red represent the South China group. b) – d) DTT analyses of male sexual traits. Shaded areas and dotted lines represent the 95% distribution and mean values of null models (based on 1,000 stochastic mappings), while solid lines represent empirical results. b) DTT analysis of male fore tibia shape. c) DTT analysis of male hind tibia shape. d) DTT analysis of male ASE shape (lateral view).

Regarding the shape data of sexual traits, the phylomorphospace analysis revealed significant divergence within clades, with considerable overlap among them. For male fore and hind tibiae, most nodes clustered near the ancestral node, while a few nodes from the EA clade were distinctly separated, suggesting the possibility of rapid phenotypic divergence in these nodes (Supplementary Fig. S14). A Mantel test among male traits found a significant correlation between the shapes of male fore tibia and hind tibia (r = 0.2831, p value = 0.001), implying potential correlated evolution. No significant correlations were detected between the shapes of male ASE and either fore tibia (r = 0.03273, p value = 0.164) or hind tibia (r = 0.08307, p value = 0.052) (Supplementary Table S7). DTT analysis identified two peaks of phenotypic divergence: male ASE diversified rapidly around 5.5 Ma, and a second peak in leg trait divergence occurred around 3.5 Ma (Fig. 4b–d). Both peaks correspond to periods of rapid speciation, though male ASE exhibited a delayed accumulation of disparity following the transition from simple to complex states (Figs. 4a–d; Supplementary Fig.13). In summary, the fore tibia and hind tibia of male exhibit a correlated evolutionary pattern in both temporal and spatial dimensions.

### Diversification Rate Estimation

Both the BSBDS and ClaDS methods identified elevated diversification rates in recently diverged taxa within each major clade, particularly in the South China group (Fig. 4a; Supplementary Fig. S15). In contrast, the basal taxa of the EA and MAL clades, *P. pusilla* and *P. gnoma*, exhibited higher diversification rates in ClaDS analysis (Supplementary Fig. S15). Additionally, most internal branches showed significantly higher net diversification rates in both methods (Fig. 4a; Supplementary Fig. S15). A notable increase in diversification rate occurred in the common ancestor of the South China group around 5 Ma (Fig. 4a). Evolutionary rate analyses of sexual conflict intensity revealed similar rate shifts in both sexes, with a marked increase in recent EA species and *P. shaanxiensis*, especially in males (Supplementary Fig. S16).

### Association Between Sexual Conflict and Speciation

Sexual conflict intensity was quantified using composite trait measures. *P. extensa*, *P. piliformis*, and *P. contorta* exhibited the strongest male-biased conflict, while *P. pusilla* showed the weakest (Fig. 4a, Supplementary Table S15). *P. contorta* also displayed the highest female conflict intensity (Fig. 4a, Supplementary Table S16). Linear regression analysis demonstrated a significant positive correlation between male and female sexual conflict, with a slope of 0.4331—indicating male-biased coevolutionary arms races (R^2^ = 0.2836, p value = 0.002). Both male and female conflict intensities were positively correlated with net diversification rates (R^2^ = 0.4153, p value = 4.107e-06 for male, and R^2^ = 0.1787, p value = 0.01784 for female). Additionally, PGLS analyses using a Brownian motion correlation structure revealed a significant positive relationship between sexual conflict intensity and net diversification rate in both sexes (t-value = 6.81, p-value < 0.001 for male; t-value = 2.61, p-value = 0.014 for female). For discrete sexual traits, species exhibiting modified male sexual conflict-related traits displayed significantly higher diversification rates than those with simpler traits (Supplementary Fig. S17). In contrast, species with elongated tergite VIII or reinforced hind femora did not show significantly higher diversification rates, and those with modified gonocoxal setae exhibited significantly lower rates (Supplementary Fig. S17).

HiSSE analysis indicated that the best-fit model for male ASE complexity was the CID-2 model (Model 6) with turnover rates 0A=0B and 1A=1B, and equal extinction fractions (Supplementary Table S8). This suggests that trait state changes were independent of diversification. In contrast, BiSSE models (Model 2), with free turnover rates and equal extinction fractions, best fitted the other three leg traits and all female traits, supporting the hypothesis that trait evolution plays a role in driving diversification (Supplementary Tables S9–S14). However, no significant increase in diversification was associated with female trait modifications in the MAL or PHI clades, as revealed by BSBDS and ClaDS analyses (Fig. 4a; Supplementary Fig. S15).

### Demographic History Inference

PSMC analysis revealed that most species across the three major clades maintained similar effective population size (*Ne*) from the Penultimate Glaciation (PG, 135–194 ka) through the Naynayxungla Glaciation (NG, 0.5–0.78 Ma), followed by declines during the NG, suggesting that *Pseudovelia* species shared a common demographic history during that period (Figs. 5c–e). Subsequently, during the Last Glacial Maximum (LGM, 19.0–26.5 ka), most MAL and PHI species experienced rapid increases in *Ne*, whereas EA species showed little change (Figs. 5c–e). This indicates that Malaysian and Philippine populations were less affected by the LGM, while East Asian populations may have been confined to small refugia during that time.

**Figure 5.**
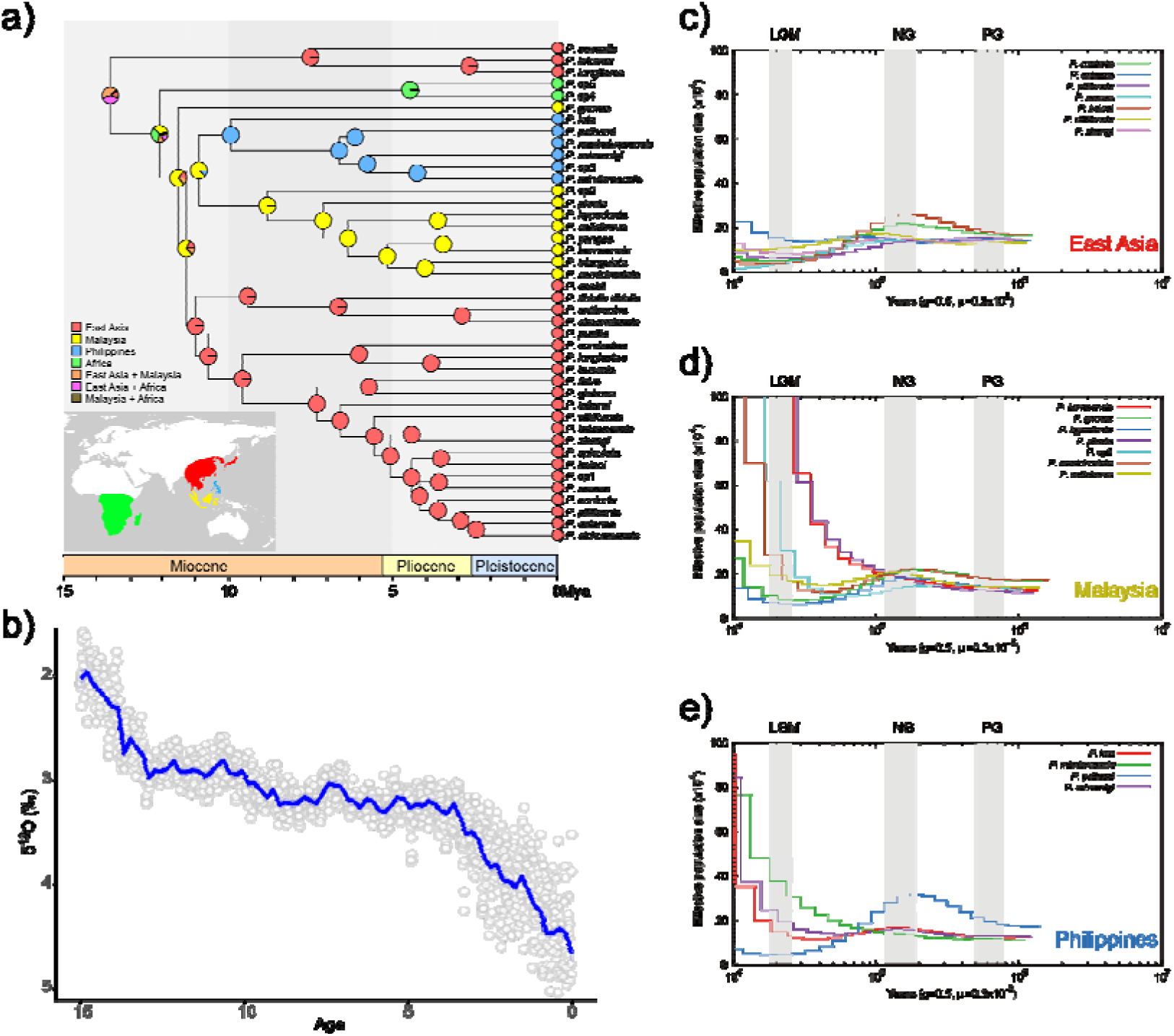
Historical biogeography and demographic history of *Pseudovelia*. a) Ancestral range estimation in BioGeoBEARS under the DEC+j model. b) Past fluctuations in global temperatures since 15 Ma, plotted from Westerhold et al. (2020). c) – e) Demographic histories of three major clades reconstructed with PSMC. Colored lines represent changes in effective population size (*N_e_*). c) East Asia clade. d) Malaysia clade. e) Philippines clade.

### Ancestral Ranges Estimation

Model comparison favored the DEC + j model over the DEC model based on log-likelihood values. Under DEC + j, the ancestral range of *Pseudovelia* was inferred as East Asia + Malaysia. The ancestor of EA+MAL+PHI clades likely originated in Malaysia, with subsequent dispersal events to East Asia and the Philippines, respectively (Fig. 5a).

## Discussion

### Introgression and ILS Leading to Wide-spread Phylogenetic Conflict

Mitochondrial paraphyly is widespread phenomenon among animals (McKay & Zink, 2010; Noguerales & Emerson, 2025; Ross, 2014). Our mitochondrial tree revealed extensive paraphyly within *Pseudoveila* species, particularly within the EA clade (Fig. 3a; Supplementary Fig. S2). This finding corroborates the previous report of mitochondrial paraphyly in *Pseudovelia* (Ye et al., 2016), and strongly challenges species delimitations based solely on morphological data. In contrast, both Maximum Likelihood (ML) and SVDquartets analyses based on SNP data recovered the monophyly of all morphologically defined species, leading support to the robustness of the existing classification (Supplementary Figs. S3 and S4). As a widely discussed topic in recent years, mito-nuclear discordance can arise from several factors, including introgression, ILS and mitochondrial capture (McKay & Zink, 2010; Noguerales & Emerson, 2025; Ross, 2014). Our analyses provide compelling evidence for both introgression and ILS within *Pseudovelia*, especially in the EA clade, identifying these processes as the primary drivers of the observed mitochondrial paraphyly.

The advent of phylogenomic approaches has greatly enhanced our ability to detect introgression and ILS (Suvorov et al., 2022; Tan et al., 2023). Using genome-scale data, we present the first comprehensive survey of these processes in *Pseudovelia*. Both introgression and ILS are known to be prevalent in rapidly radiating clades, significantly influencing species tree inference. While concatenated methods have traditionally been used for estimating species trees, they may yield inaccurate phylogenetic estimates, particularly under high levels of introgression or ILS (Mirarab et al., 2014; Roch & Steel, 2015). In contrast, the Multi-species Coalescent (MSC) method can improve the accuracy of phylogenetic estimation and is particularly well-suited for addressing the incongruence between gene trees and species trees resulting from introgression or ILS (Jiang et al., 2020; Wascher & Kubatko, 2021). In our study, the concatenated methods produced unstable topologies among species within the South China group, which contrasted with the topology recovered by the MSC species tree (Supplementary Figs. S3–S6). In contrast, the MSC species trees, based on both SNP sliding windows and USCOs, exhibited relatively congruent topologies within the South China group (Supplementary Figs. S5 and S6). Moreover, analyses using QuIBL, f-branch, and Phytop revealed a stronger signal of both introgression and ILS within the South China group. These findings demonstrate that introgression and ILS have significantly influenced the phylogenetic inference of *Pseudovelia* especially within South China group, and the MSC method can mitigate their confounding effects to some extent.

### Phylogenetic Heterogeneity and Asymmetric Sexual Conflict in Pseudovelia

Sexual conflict traits are known to exhibit mosaic evolution, characterized by clade-specific evolutionary patterns and notable asymmetry between sexes (Carvalho et al., 2021; Crumière et al., 2019; Genevcius et al., 2020; Miller, 2003). In our study system, males have independently evolved multiple novel tibial traits exclusively within the EA clade, whereas putative female counter-adaptations are clade-specific and non-overlapping: an elongated tergite VIII in the MAL clade, bristle-like setae on the gonocoxa in the PHI clade, and strengthened hind femora in the EA clade (Fig. 4a; Supplementary Fig. S13). This distribution indicates a clade-specific pattern of sexual trait evolution. Such mosaic evolution likely reflects deep isolation and divergent evolutionary pressures across major clades. This phylogenetic heterogeneity suggests that historical isolation and divergent coevolutionary pressures across major clades have played a significant role in shaping sexual conflict. These pressures may stem from early divergence and radiation in geographically isolated regions subjected to distinct climatic fluctuations. Over time, limited gene flow facilitated the independent emergence of clade-specific sexual conflict dynamics, with varying ecological conditions further influencing the fitness of sexual traits and driving divergent coevolutionary trajectories.

Within the EA clade, males have undergone a burst of diversification in grasping structures, whereas females have shown no structural modifications comparable to those in the MAL or PHI clades, instead only developing strengthened hind femora in a few species. We propose two potential mechanisms that could explain this asymmetry: 1) Female resistance failure due to fitness costs: Females may fail to evolve resistance traits if the fitness cost of resisting males exceeds the cost of multiple mating (Gavrilets et al., 2001). While this hypothesis is theoretically plausible, it is difficult to empirically validate in our system, particularly given the difficulty in directly measuring the female cost. Additionally, females may exhibit cryptic or undefined resistance traits, making the analysis of female cost challenging. 2) Asymmetric coevolution: Coevolution may not be perfectly balanced, leading to a situation where one sex gains an advantage over the other during the coevolutionary process (Arnqvist & Rowe, 2002). We propose that this hypothesis aligns more closely with our data for several reasons. As a rapidly radiated lineage, male sexual conflict traits underwent recent, explosive diversification, and our DTT analysis confirmed that key male conflict traits evolved rapidly during the final stages of lineage radiation (Figs. 4b and c). Meanwhile, female counter-adaptations may have evolved more slowly, resulting in a transient male advantage. The time lag in sexual trait evolution has also been observed in Coleoptera (Genevcius et al., 2020; Miller, 2003). Importantly, East Asia cladebecause these two hypotheses are not mutually exclusive, we suggest that both cost imbalances and transient asymmetries in coevolutionary trajectories could operate synergistically to shape the observed male-biased sexual conflict in *Pseudovelia*.

### Integrated “Trait package” of Male Grasping Traits as a Driver of Speciation

Most studies have confirmed that sexual conflict is a driving factor of speciation (Arnqvist et al., 2000; Gavrilets, 2000, 2014), although some studies have presented opposing conclusions (Carvalho et al., 2021). *Pseudovelia* represents an ideal model system to investigate this relationship due to its high species diversity and extensive variation in male sexual traits. By integrating multiple traits, we quantified the intensity of sexual conflict across species and found a significant positive correlation with net diversification rates. Additionally, species possessing these putative male sexual conflict-related morphological modifications exhibited significantly higher net diversification rates compared to those lacking such modifications (Supplementary Fig. S17). HiSSE (Hidden State Speciation and Extinction) analyses further support the hypothesis that evolutionary changes in three key male sexual conflict traits are associated with increased diversification (Supplementary Tables S9–11). These findings align with the theory that sexual conflict drives speciation (Arnqvist et al., 2000; Gavrilets, 2000, 2014), particularly when traits work synergistically to enhance male grasping ability.

Functional and evolutionary analyses reveal that leg modifications in EA males evolved recently and may serve to immobilize resistant females during initial grasping (Fig. 1a), thereby indicating intensified sexual conflict. Importantly, these traits form a functional “trait package” that collectively improves grasping ability and exacerbates sexual conflict. Mantel tests revealed a significant correlation between the shapes of the foreleg and hind leg (Supplementary Table S7), suggesting potential correlated evolution. HiSSE analyses confirm that these traits are significant drivers of diversification (Supplementary Tables S9–S11), supporting the view that coordinated evolution of coercive traits promotes speciation. This contrasts with findings in Acraeini butterflies, where a post-copulatory trait (sphragis) showed no association with diversification (Carvalho et al., 2021). The discrepancy likely stems from the functional differences between the traits: in *Pseudovelia*, traits enhance pre-copulatory coercion, whereas the sphragis functions as a passive mating plug without contributing to male mating success.

### Low-intensity Conflict allowing Evolutionary Flexibility in Female Resistance

During copulation, males first grasp females with their legs before utilizing ASE to open the female proctiger, suggesting that male ASE typically functions after a successful grasp has been achieved (Figs. 1a and b). Ancestral state reconstruction indicates that complex male ASE was already present in the common ancestor of the three major clades, and phenotypic divergence in ASE preceded that of leg grasping traits, as supported by DTT analyses (Figs. 4a–d; Supplementary Fig. 13). In contrast, the male leg modifications evolved much later and are restricted to recently diverged lineages within East Asia clade. We therefore propose that the absence of male modified leg structures imposed only moderate antagonistic pressure on females, thereby maintaining sexual conflict at a relatively low intensity during the early phase of ASE evolution. This moderate pressure allowed for evolutionary flexibility in putative female resistance strategies, led to lineage-specific counter-adaptations among females, generating a mosaic pattern of traits across subclades.

Furthermore, we propose that highly divergent ASE contributed to prezygotic isolation, facilitating species coexistence within geographically restricted areas and helping to maintain genetic diversity, which later promoted diversification events. Under this scenario, the emergence of complex ASE initially arose through sexually antagonistic evolution, but their subsequent diversification may have been maintained by prezygotic isolation. This role of genital diversification in prezygotic isolation has been observed in other radiations (Tu et al., 2025). Thus, while ASE divergence did not directly promote speciation, its contribution to reducing gene flow and sustaining diversity under conditions of low-intensity conflict likely supported subsequent lineage-specific evolutionary trajectories.

Finally, we emphasize that the grasping and anti-grasping functions of the male and female traits examined here have been inferred primarily from morphological analogies with Gerridae, rather than experimentally demonstrated in *Pseudovelia*. Future functional experiments, such as mating trials with trait-manipulated individuals, or high-speed video analysis of copulatory interactions, are needed to directly quantify the mechanical roles of these structures and their fitness consequences for both sexes. Integrative approaches combining behavioral experiments with phylogenomic and ecological analyses will be essential for understanding the relative contributions of sexual conflict, reproductive interference, and environmental change in shaping the diversification of this lineage.

### Why the South China Group Became a Diversity Hotspot

Our study revealed that the South China group within the EA clade represents a critical nexus for both speciation and trait diversification in *Pseudovelia*. This lineage exhibits the highest net diversification rate and phenotypic disparity, particularly in traits related to sexual conflict. Key innovations are concentrated within the South China group. The fore tibial process and spine-like setae on the hind tibiae evolved exclusively within the EA clade, with a single-origin innovation in the South China group. In contrast, curved hind tibiae evolved independently twice within the EA clade, i.e., once in the South China group and once in *P. shaanxiensis* (Fig. 4a; Supplementary Fig. S13). This spatial and temporal clustering of novel traits suggests that the South China group serves as a primary center of diversification in *Pseudovelia*. Moreover, our BAMM analysis revealed a significant increase in the evolutionary rate of sexual conflict intensity in both males and females, implying an escalating arms race between the sexes (Supplementary Fig. S16). We propose three synergistic factors driving this pattern: 1) Sexual conflict as a catalyst for speciation: Early diversification of male ASE established a baseline for reproductive isolation through mechanical matches. Subsequent elaboration of tibial grasping structures heightened physical stress during pre-mating struggles. The correlated evolution of this “trait package” likely intensified sexual conflict, promoting speciation through arms races. 2) Genomic turbulence within the South China group: Phylogenomic analyses revealed exceptionally high levels of introgression and ILS in the South China group compared to other *Pseudovelia* lineages. These processes may have preserved critical genetic variation for trait innovation and/or directly facilitated speciation via adaptive introgression. 3) Geographic and climatic drivers: The South China group radiated within the terrestrial “sky islands” of South and Southeast China. Geographic isolation, in combination with climatic fluctuations, facilitated speciation in this region (Ye et al., 2016). Our results also indicate that the EA clade was more strongly impacted by the Neogene and Last Glacial Maximum than other lineages. Additionally, global temperatures have steadily declined since 5 Ma (Fig. 5b), coinciding with the diversification of the South China group. The climate oscillations during this period may have triggered cycles of isolation and connection among sky islands, promoting speciation by allowing recombination of adaptive variants accumulated during isolation, consistent with the Mixing-Isolation-Mixing model (He et al., 2019; Seehausen, 2004). The interaction of these evolutionary, genomic, and environmental forces created ideal conditions for the rapid radiation of the South China group.

## Supplementary Material

Supplementary data are available at Molecular Biology and Evolution online.

## Acknowledgments

This project is funded by the National Natural Science Foundation of China (No. 32322012, No. 32470467), the Natural Science Foundation of Tianjin, China (24JCYBJC01910), and the Fundamental Research Funds for the Central Universities, Nankai University (No. 92612038).

## Author Contributions

The study was conceived by Z.Y. and W.J.B., and experiments were designed by Z.H.L., H.Y.C., Z.Z.J., Z.Y. and W.J.B. Sample collection was carried out by Z.H.L., Z.Y., Z.Z.J., H.F., C.H., H.Z., S.Y.F., C.L. and M.Q. The data were analyzed by Z.H.L., H.Y.C., Z.Z.J. and B.X.G.. The article was written by Z.H.L. and Z.Y.

## Data Availability Statement

Our resequencing data were deposited at GenBank (https://www.ncbi.nlm.nih.gov/bioproject/PRJNA1327476/).

