## Supplementary Figures for "Phylogenetic Mosaic of an Arms Race with Asymmetrical Sexual Conflict and Its Macroevolutionary Consequences in a Lineage of Small Water Striders"

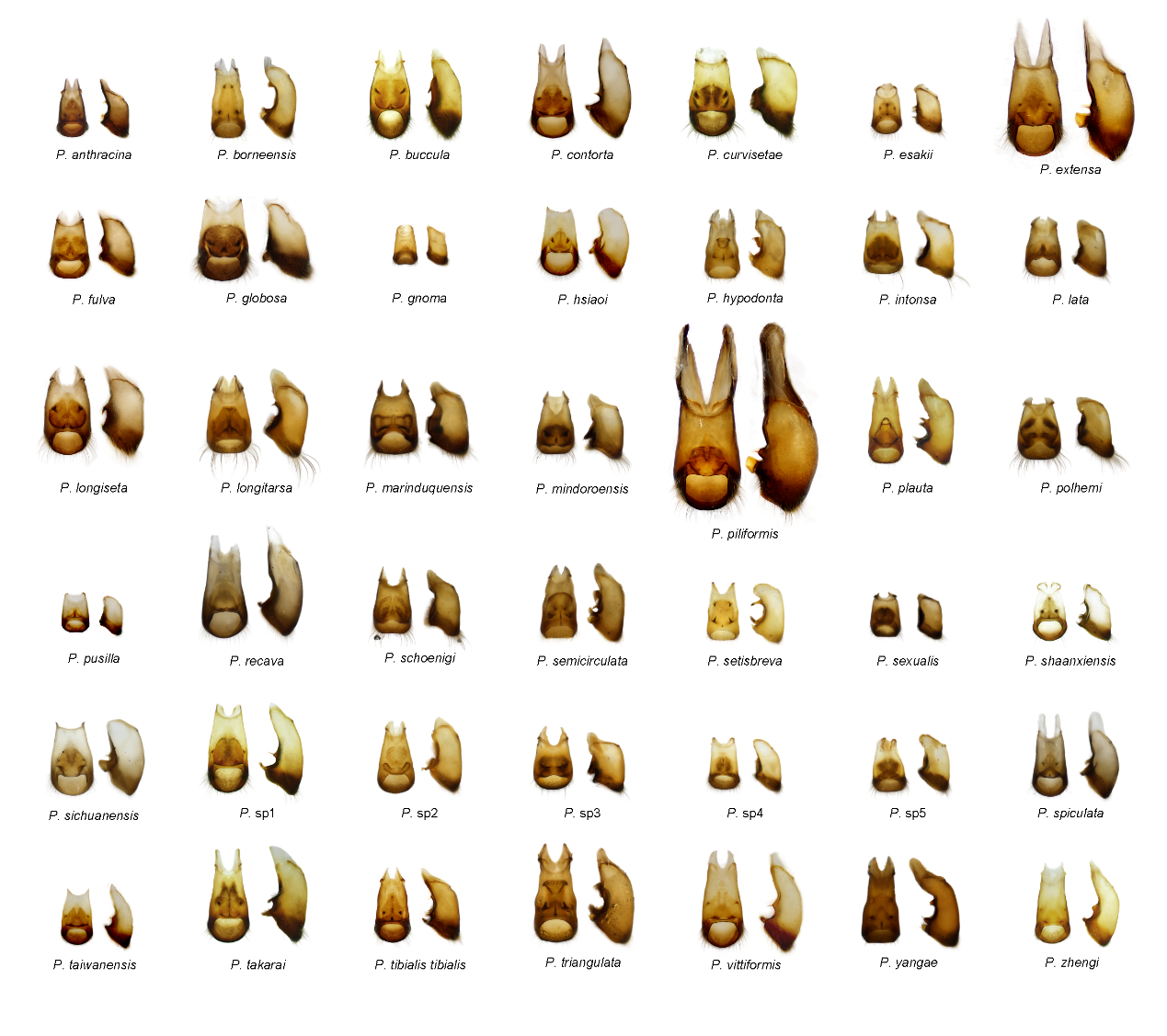

**Figure S1. Morphological disparity of the male ASE of *Pseudovelia*.** Photographs show the ventral and lateral views of the male ASE.

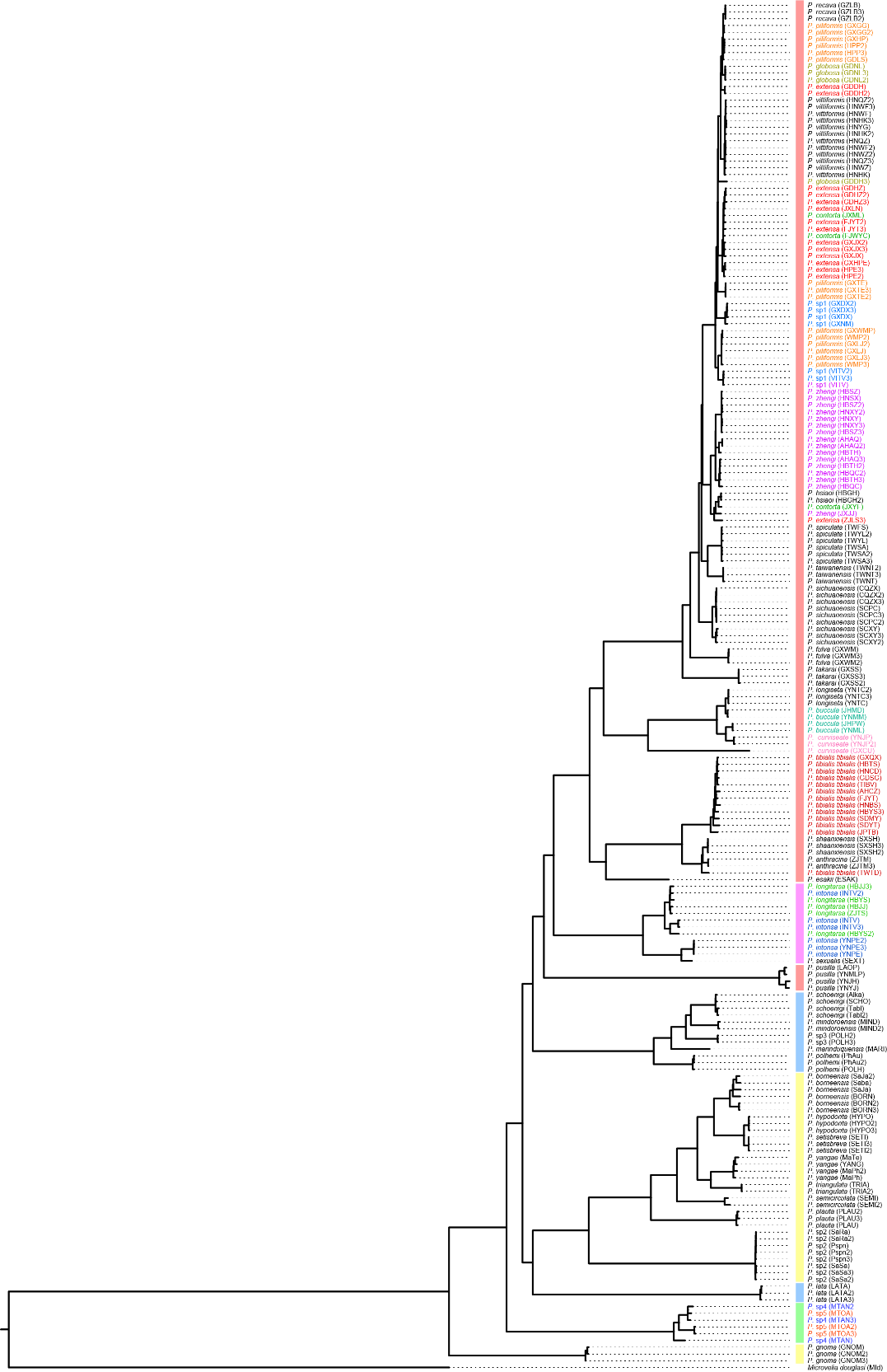

**Figure S2. Concatenated tree inferred using the maximum likelihood (ML) method in IQ-tree2 based on the mitochondrial genome dataset.** Colored species represent non-monophyly taxa.

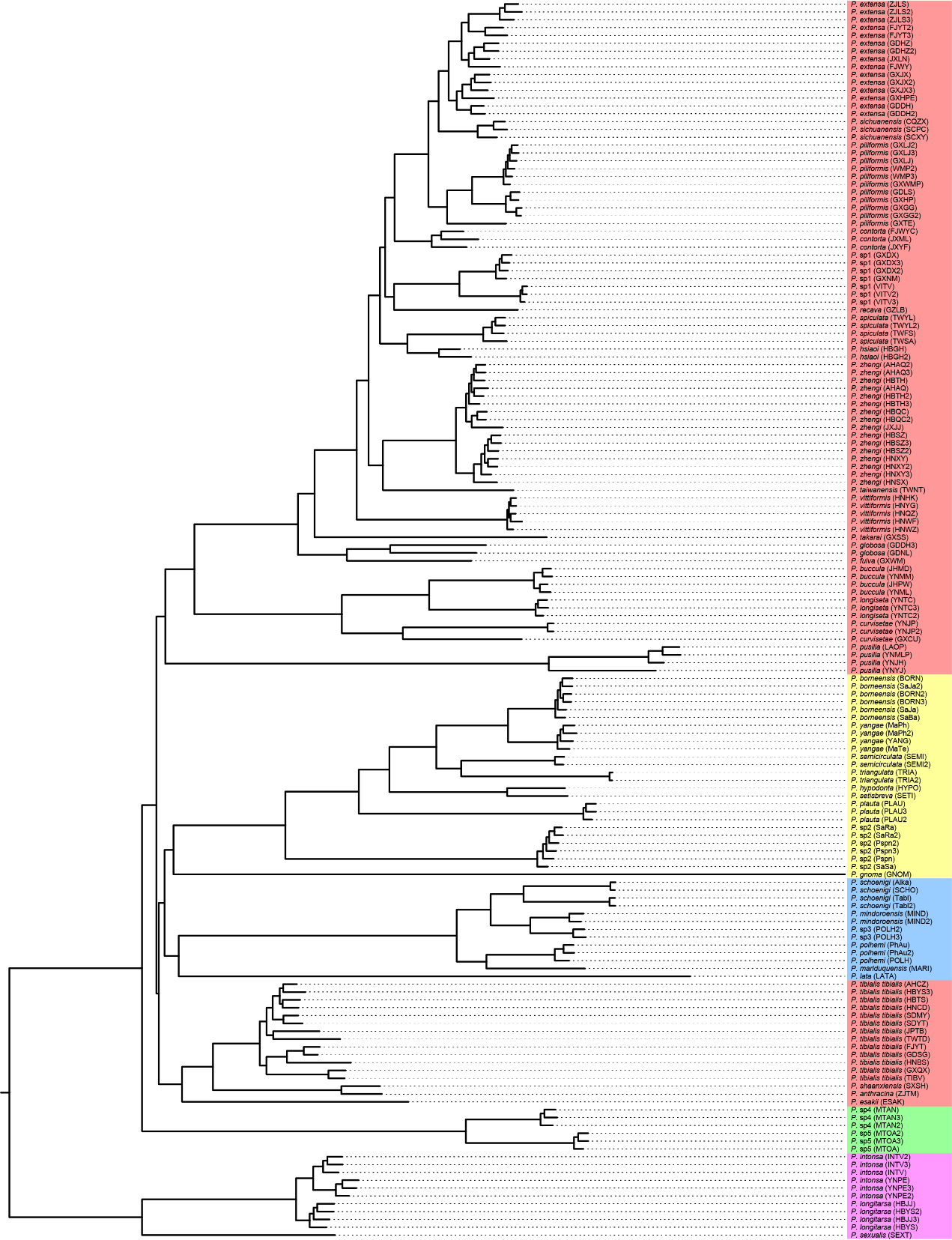

**Figure S3. Concatenated tree inferred using the ML method in IQ-tree2 based on the concatenated SNP dataset.**

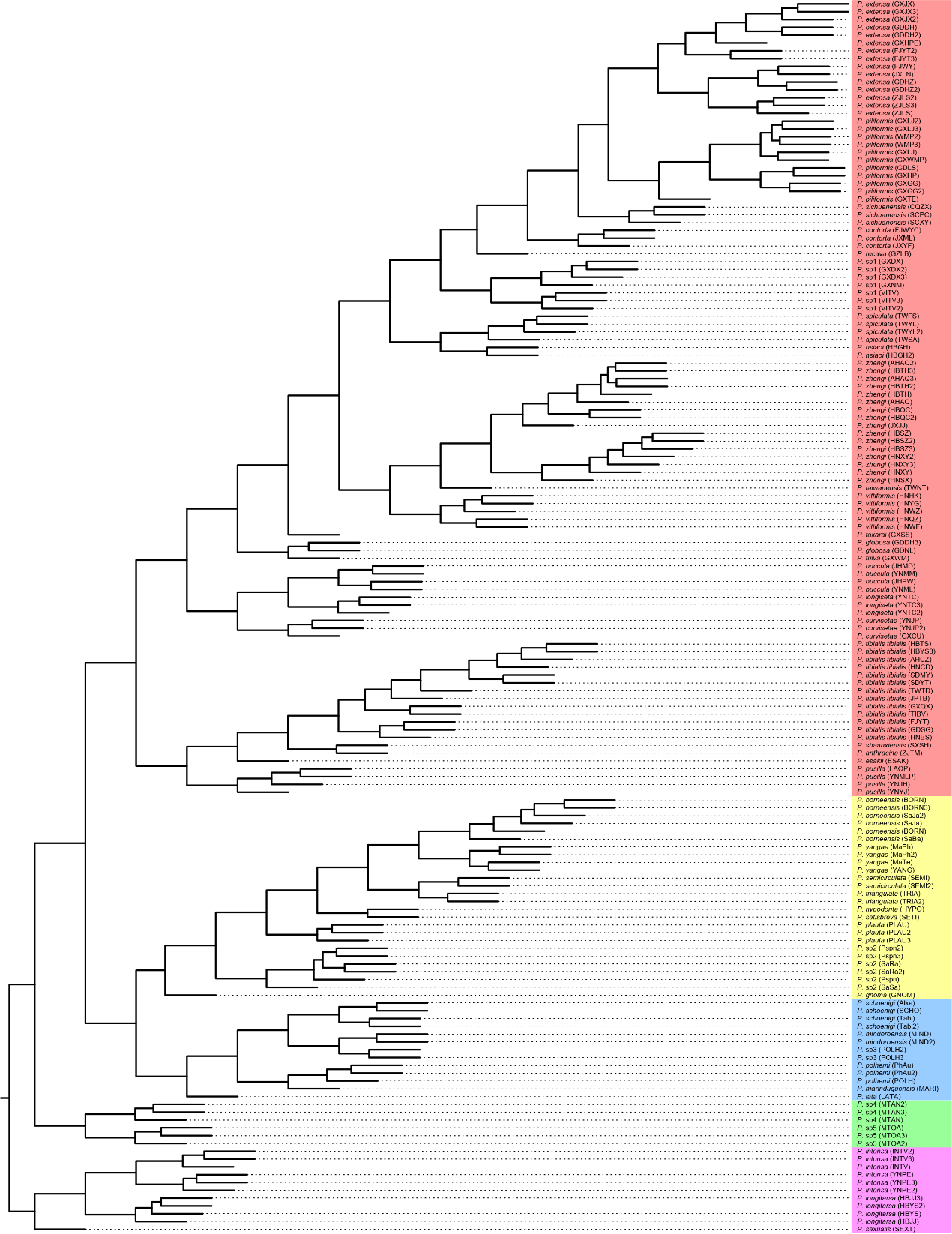

**Figure S4. Coalescent tree inferred under the multispecies coalescent (MSC) model using the SVDquartets algorithm in PAUP* based on the concatenated SNP dataset.**

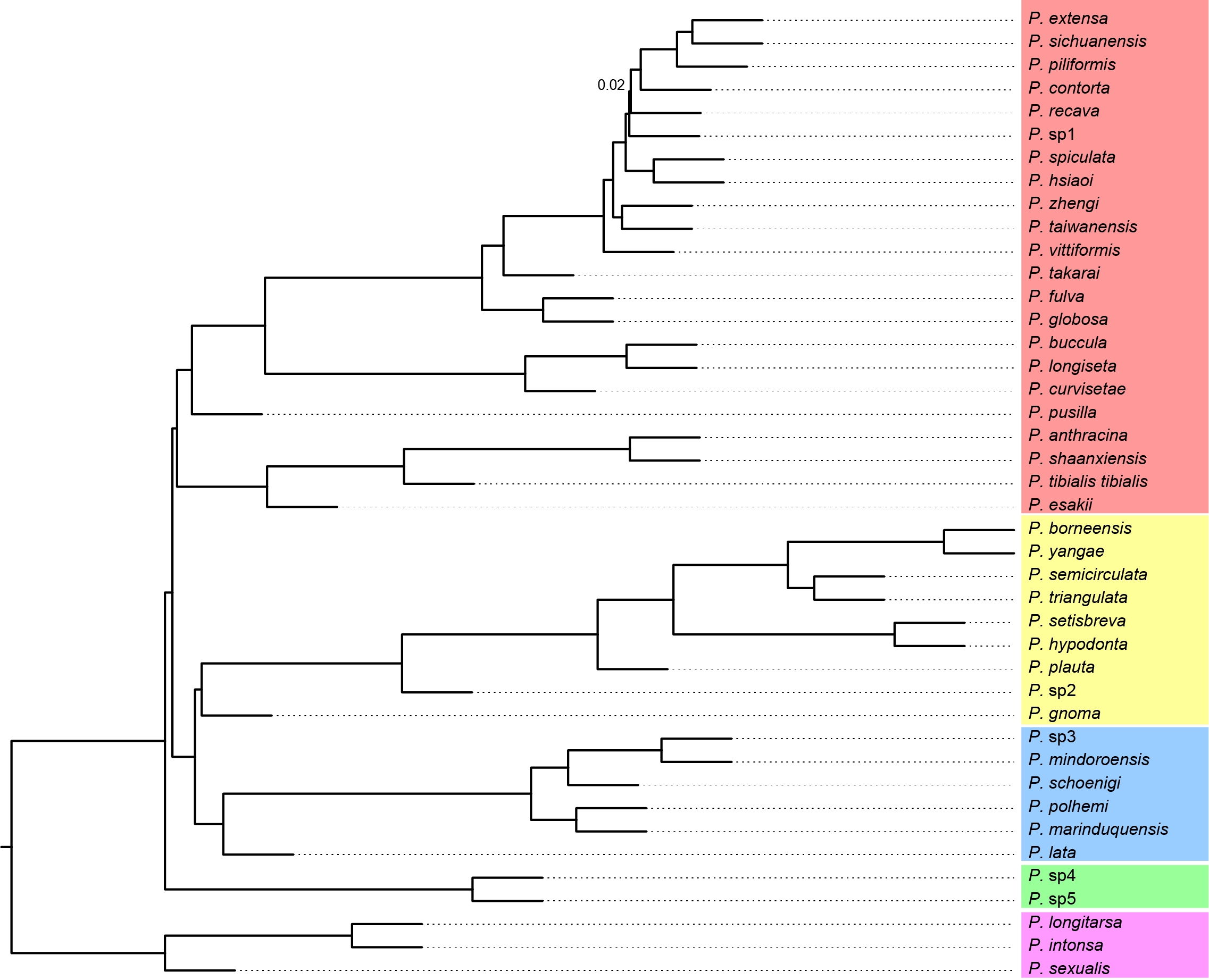

**Figure S5. Species tree inferred under the MSC model using ASTRAL based on SNP sliding windows.** The phylogenetic tree was constructed using whole genome SNPs divided into 20 kb windows with a 400 kb step length. Numbers on the nodes represent support values; nodes without numbers are 100% supported.

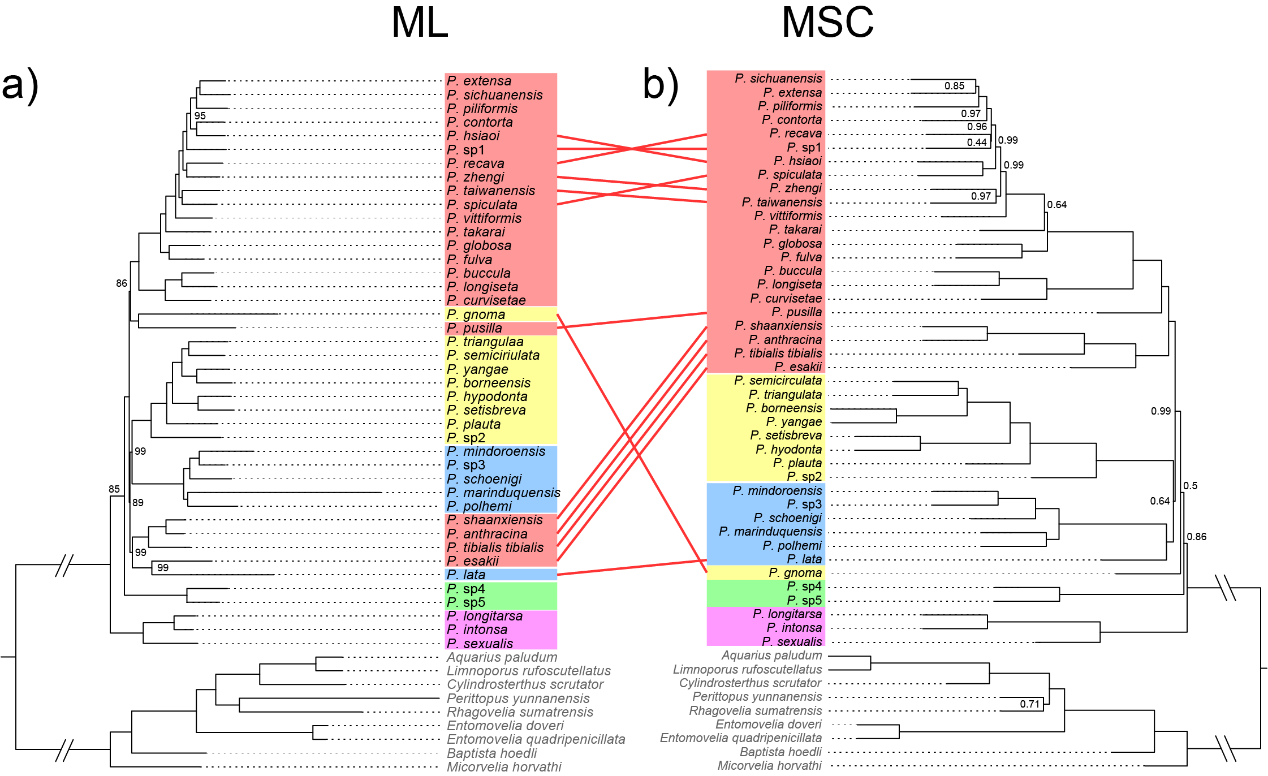

**Figure S6. Comparison of topologies inferred from the ML method (left) and the MSC method (right) based on the 75% completeness USCOs dataset.** The ML tree was reconstructed using IQ-tree2, and the MSC tree was reconstructed using ASTRAL. Numbers on the nodes indicate support values; nodes without numbers are 100% supported.

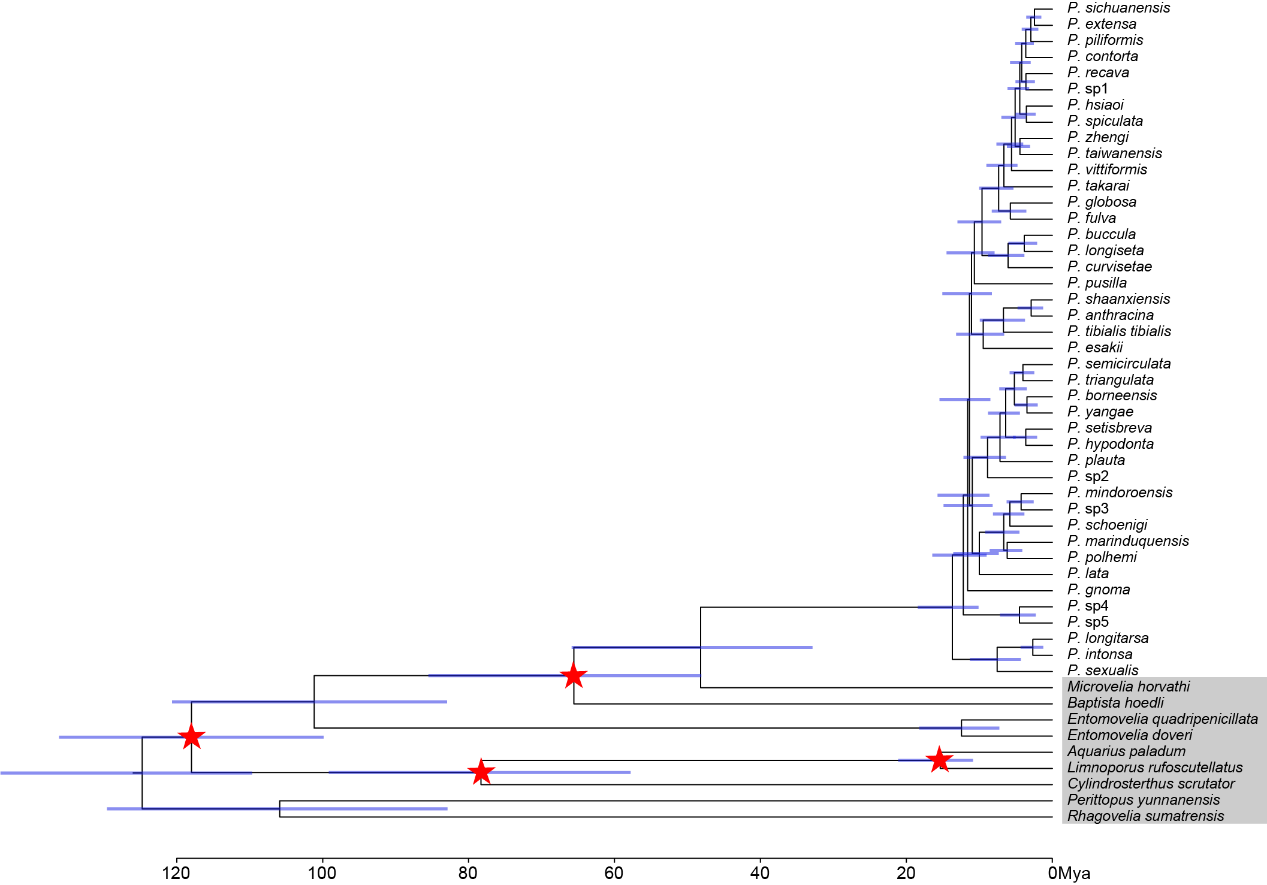

**Figure S7. Divergence time estimation of *Pseudovelia*.** The estimation was based on the 75% completeness USCOs matrix and the topology of the MSC tree inferred using ASTRAL. Analyses were conducted with MCMCtree in PAML. Node bars represent the 95% highest posterior density (HPD) intervals of the estimated times. Red stars indicate fossil calibration nodes.

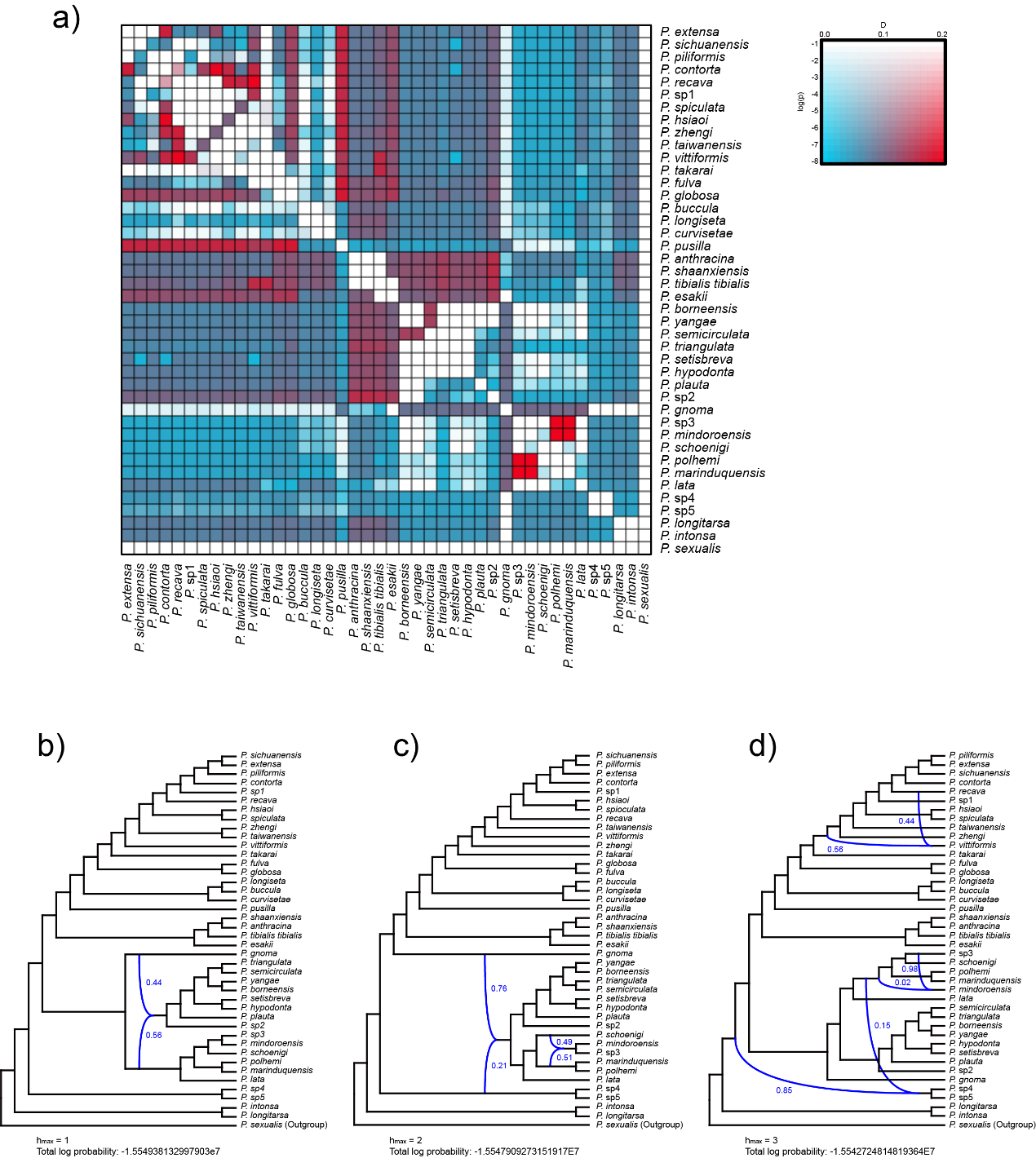

**Figure S8. Introgression and phylogenetic network analyses of *Pseudovelia*, with *P. sexualis* as the outgroup.** a) Matrix of D-statistic values from Dsuite analysis. Darker colors indicate smaller p-value, while red indicates higher D-statistic values compared to blue. b) – d) Phylogenetic network analysis in PhyloNet with h set from 1 to 3, Blue lines represent reticulation events, and numbers indicate inheritance probabilities. b) h = 1. c) h = 2. d) h = 3.

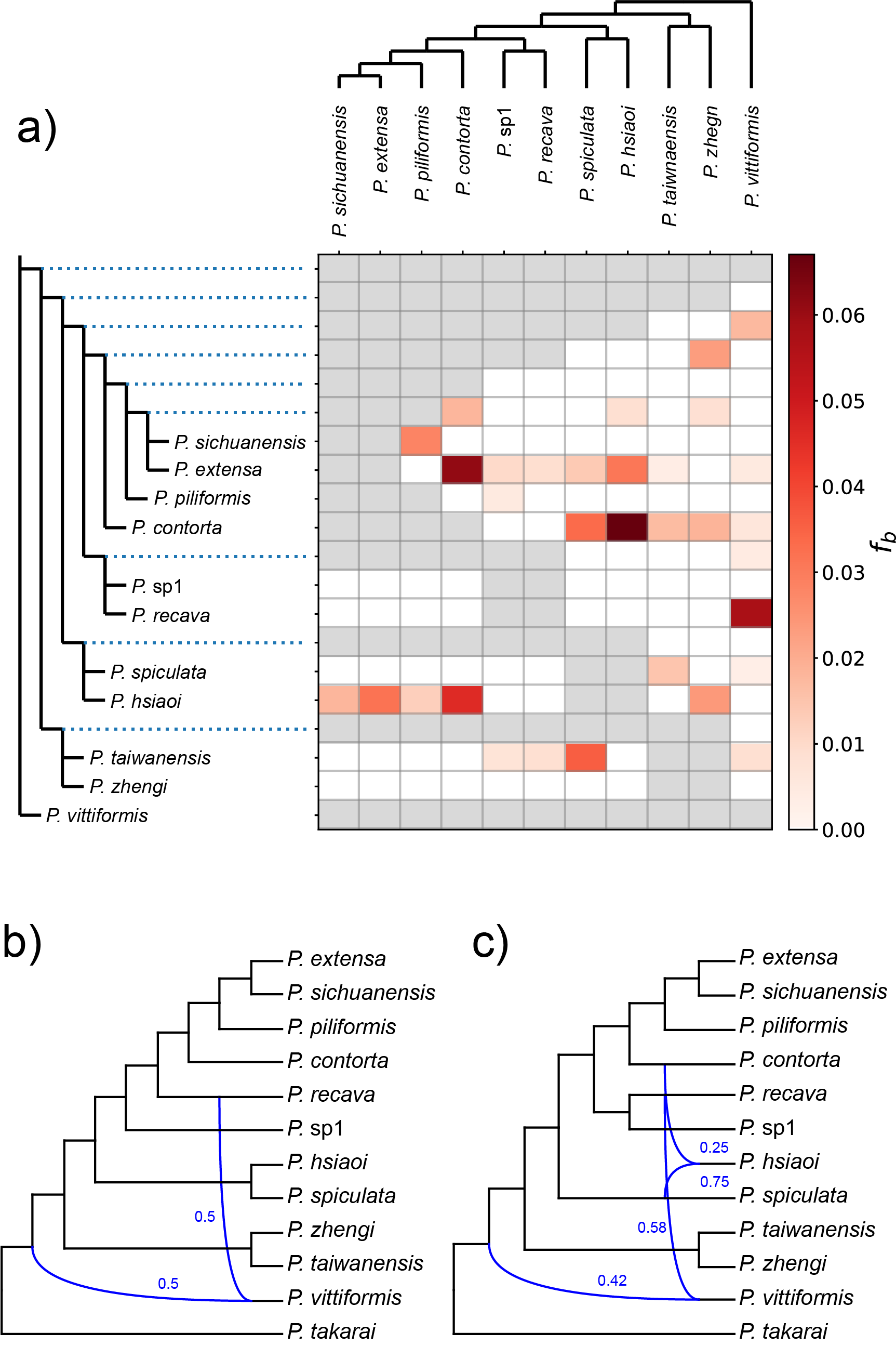

**Figure S9. Introgression and phylogenetic network analyses of the SC group.** a) Matrix of F-branch values from Dsuite analysis. b) – c) Phylogenetic network analysis in PhyloNet with h set to 1 and 2, Blue lines represent reticulation events, and numbers indicate inheritance probabilities. b) h = 1. c) h = 2.

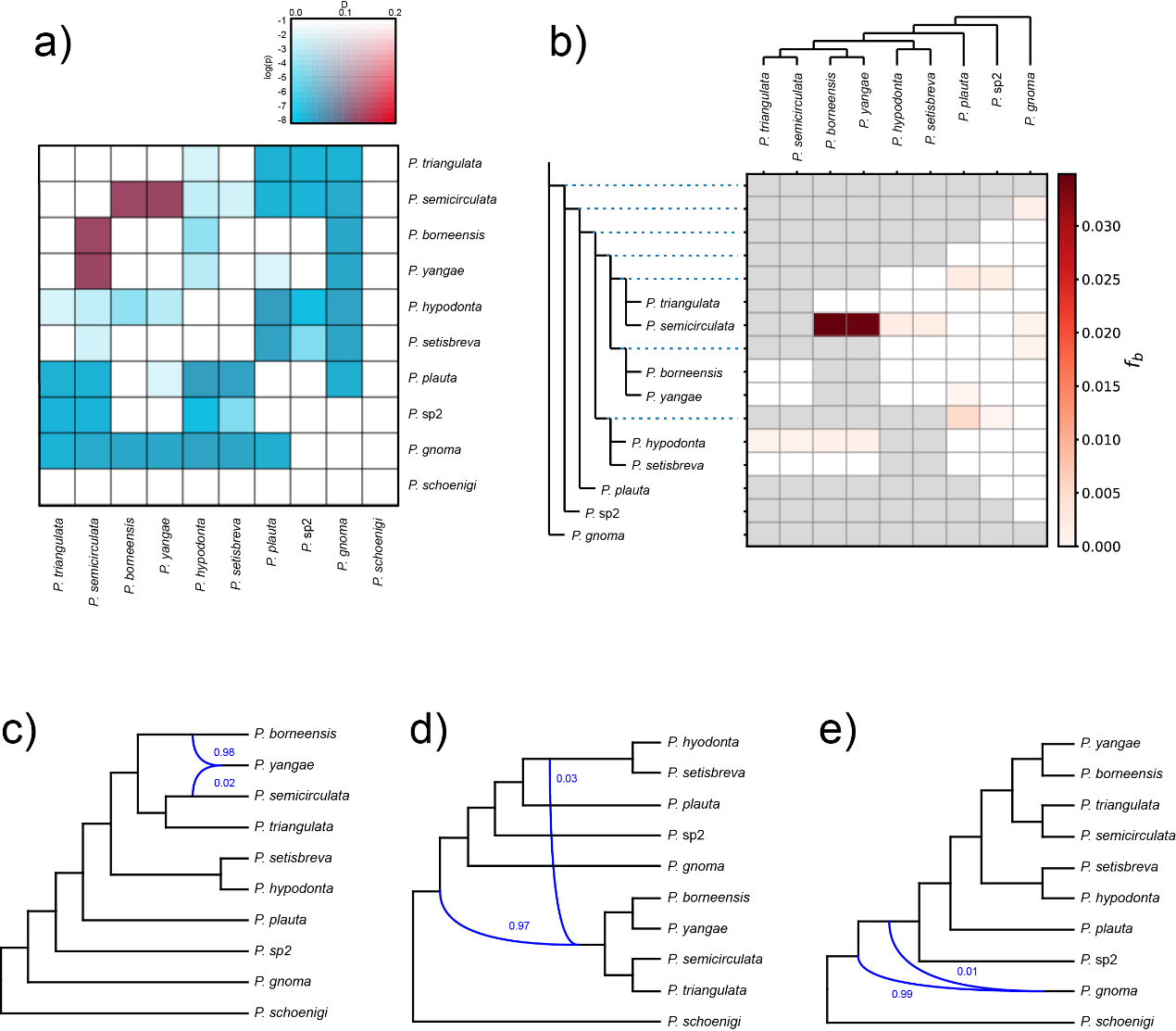

**Figure S10. Introgression and phylogenetic network analyses of the MAL clade.** a) Matrix of D-statistic values from Dsuite analysis. b) Matrix of F-branch values from Dsuite analysis. c) – e) Phylogenetic network analysis in PhyloNet with h set from 1 to 3. c) h = 1. d) h = 2. e) h = 3.

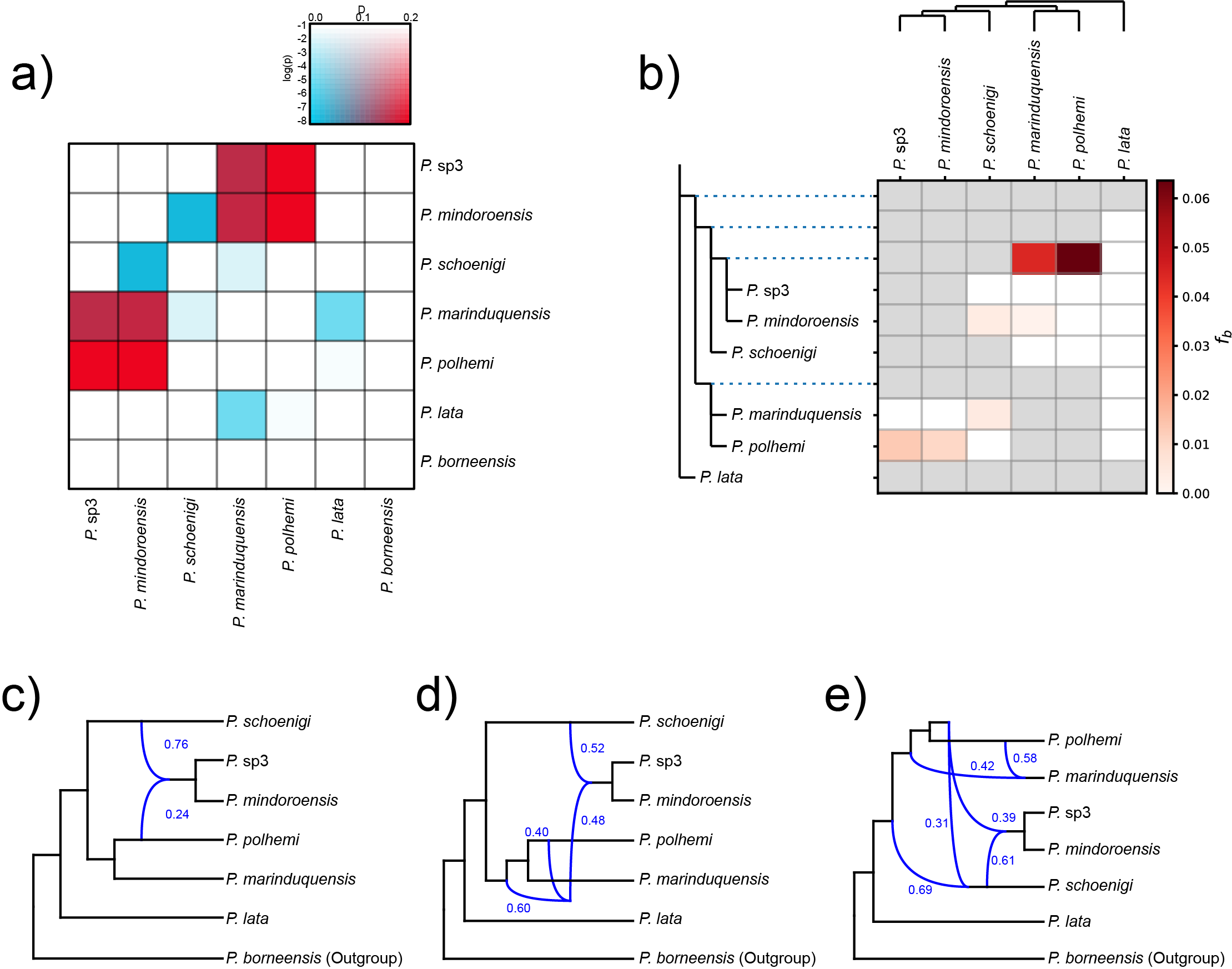

**Figure S11. Introgression and phylogenetic network analyses of the PHI clade.** a) Matrix of D-statistic values from Dsuite analysis. b) Matrix of F-branch values from Dsuite analysis. c) – e) Phylogenetic network analysis in PhyloNet with h set from 1 to 3. c) h = 1. d) h = 2. e) h = 3.

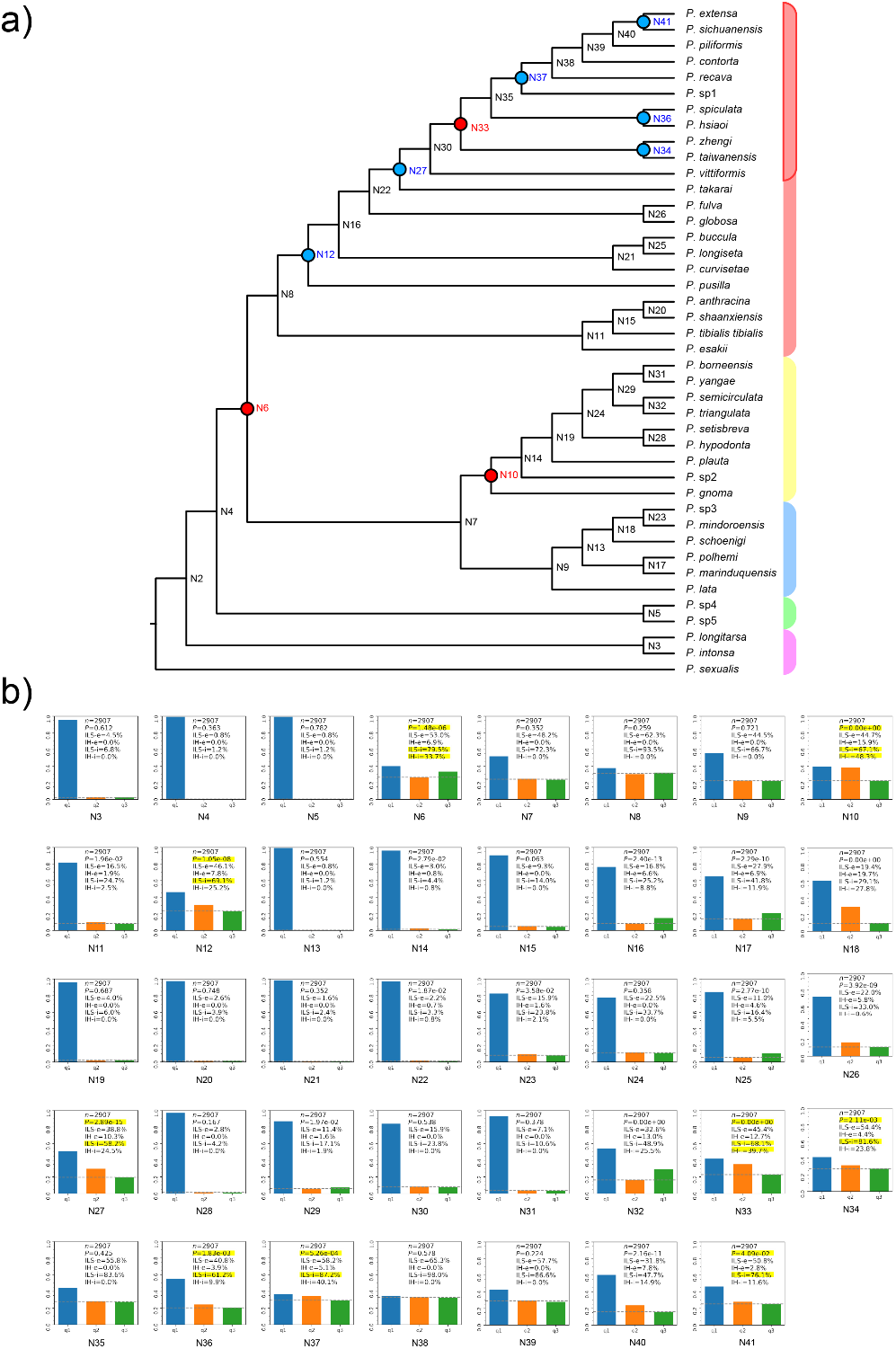

**Figure S12. Introgression and incomplete lineage sorting (ILS) analyses of *Pseudovelia* in Phytop based on SNPs sliding window trees.** a) Internal nodes showing introgression and ILS signals. Nodes in red indicate significant introgression signals only, while nodes in blue indicate both significant introgression and ILS signals. b) Intensity of introgression and ILS across nodes.

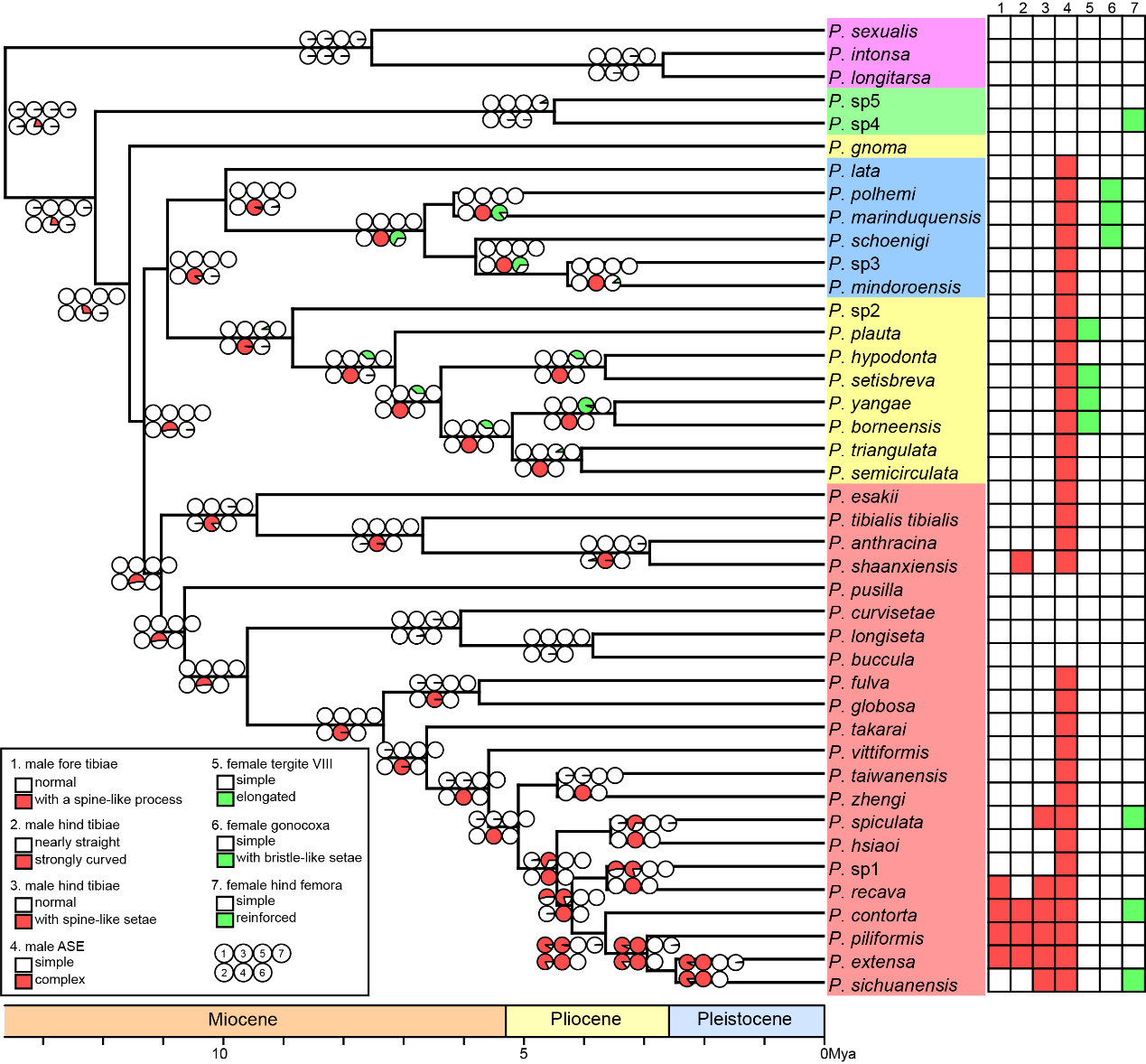

**Figure S13. Ancestral state reconstruction of sexual traits.** Male traits are colored in red and female traits in green. Colored squares represent the state of each species, while circles on internal nodes represent ancestral states estimated from 1,000 stochastic mappings in phytools.

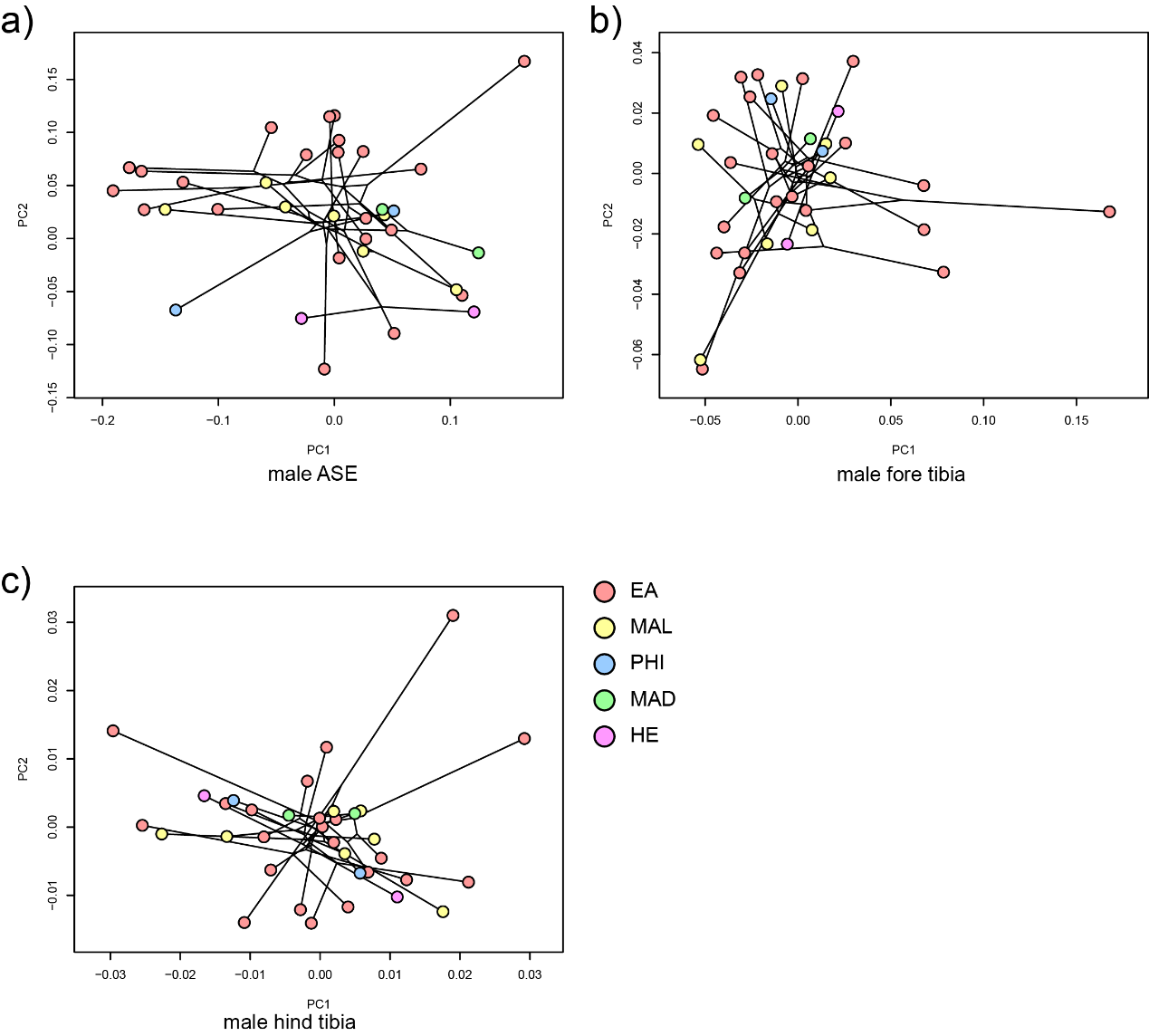

**Figure S14. Phylomorphospace of sexual traits based on shape data, grouped by major clades**. a) Male abdominal segment VIII (ASE) in lateral view. b) Inner margin of the male fore tibia. c) Inner margin of the male hind tibia.

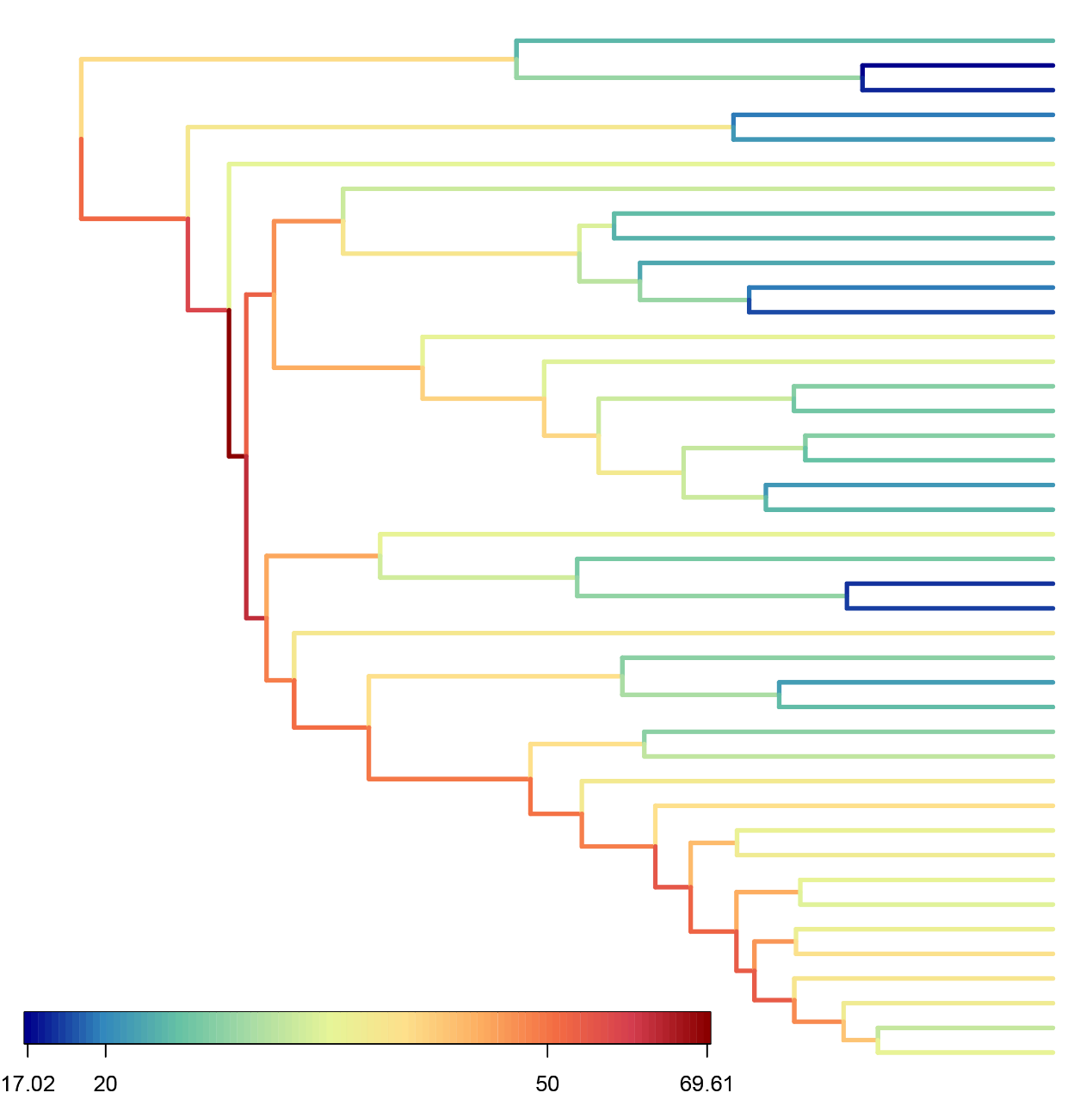

**Figure S15. Net diversification rate estimated using ClaDS in PANDA.** The phylogenetic tree is colored according to branch-specific net diversification rates.

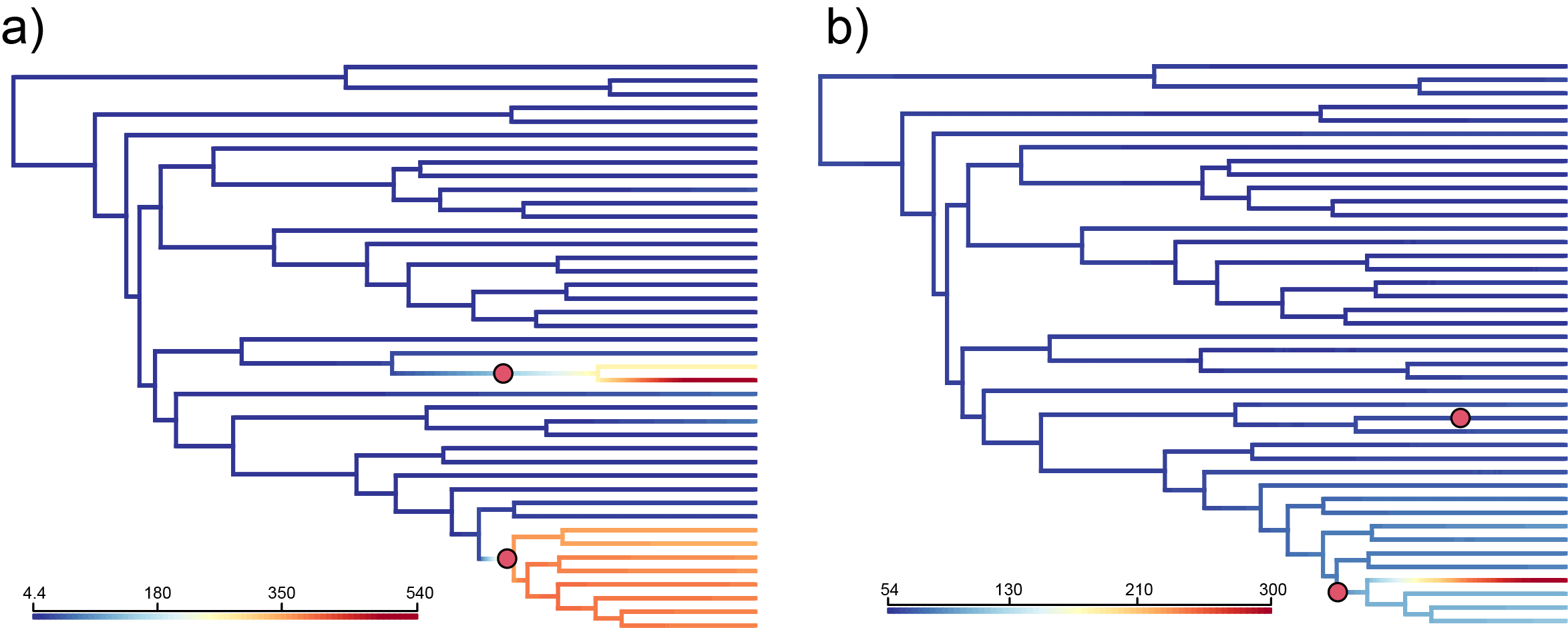

**Figure S16. Dynamics of trait evolutionary rates across the phylogeny inferred using BAMM.** Branches are colored according to their inferred evolutionary rates. Red dots on nodes indicate significant shifts in evolutionary rate. a) Rates inferred based on the intensity of male sexual conflict. b) Rates inferred based on the intensity of female sexual conflict.

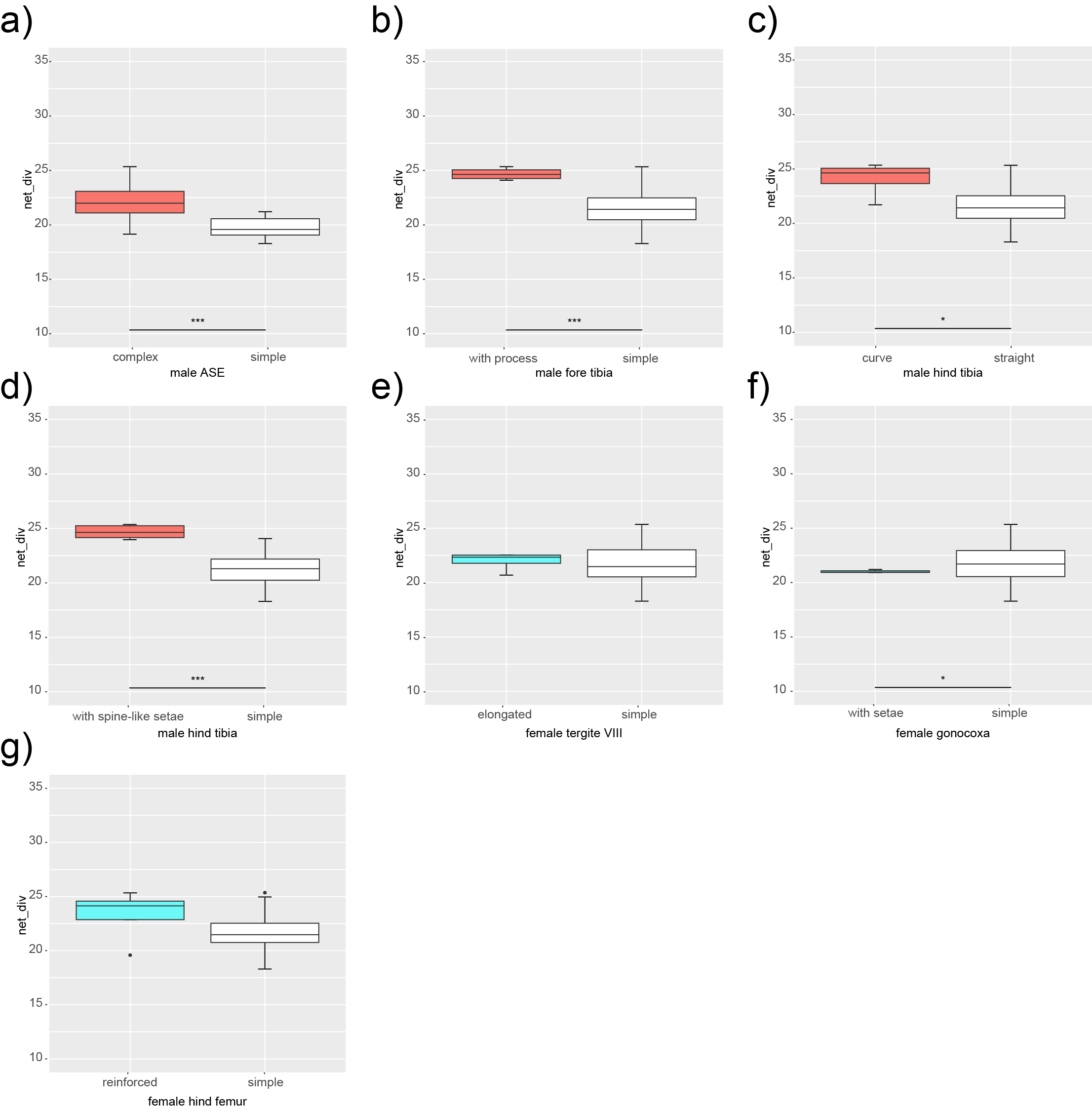

**Figure S17. Boxplots of the distribution of net diversification rates across states of sexual traits.** Rates were extracted from Revbayes BSBDS analysis. Significance was assessed using an independent two-sample t-test. a) complexity of the male ASE. b) Presence of a fore tibial spine-like process. c) Curvature of the male hind tibia. d) Presence of spine-like setae on the male hind tibia. e) Length of female tergite VIII. f) Presence of modified setae on the female gonocoxa. g) Strength of the female hind femur.
